# Calcium-binding protein S100A6 interaction with VEGF receptors integrates signaling and trafficking pathways

**DOI:** 10.1101/2021.07.29.454311

**Authors:** Leyuan Bao, Gareth W. Fearnley, Chi-Chuan Lin, Adam F. Odell, Ana C. Redondo, Gemma K. Kinsella, John B. C. Findlay, John E. Ladbury, Michael A. Harrison, Sreenivasan Ponnambalam

## Abstract

The mammalian endothelium which lines all blood vessels responds to soluble factors which control vascular development and sprouting. Endothelial cells bind to vascular endothelial growth factor A via two different receptor tyrosine kinases (VEGFR1, VEGFR2) which regulate such cellular responses. The integration of VEGFR signal transduction and membrane trafficking is not well understood. Here, we used a yeast-based membrane protein screen to identify VEGFR-interacting factor(s) which modulate endothelial cell function. By screening a human endothelial cDNA library, we identified a calcium-binding protein, S100A6, which can interact with either VEGFR. We found that S100A6 binds in a calcium-dependent manner to either VEGFR1 or VEGFR2. S100A6 binding was mapped to the VEGFR2 tyrosine kinase domain. Depletion of S100A6 impacts on VEGF-A-regulated signaling through the canonical mitogen-activated protein kinase (MAPK) pathway. Furthermore, S100A6 depletion caused contrasting effects on biosynthetic VEGFR delivery to the plasma membrane. Co-distribution of S100A6 and VEGFRs on tubular profiles suggest the presence of transport carriers that facilitate VEGFR trafficking. We propose a mechanism whereby S100A6 acts as a calcium-regulated switch which facilitates biosynthetic VEGFR trafficking from the TGN-to-plasma membrane. VEGFR-S100A6 interactions thus enable integration of signaling and trafficking pathways in controlling the endothelial response to VEGF-A.

## Introduction

Receptor tyrosine kinases (RTKs) are integral membrane proteins and enzymes which regulate essential features of cell, organ, tissue and animal function (Lemmon et al., 2016). RTK binding to exogenous ligands enables the transmission of signals into the cell interior through activation of multiple signal transduction pathways. In spite of numerous studies on different RTKs over the past 50 yrs, we still lack an understanding of how different cells integrate RTK activation, signaling and cellular responses. This is further complicated by discovery that post-translational modifications such as phosphorylation and ubiquitination can also modulate trafficking and proteolysis. This is important as the presence of activated RTK complexes at different intracellular locations could activate different signal transduction pathways.

An archetypal RTK is a Type I membrane protein with a glycosylated, extracellular N-terminus which is used to ‘sense’ exogenous soluble and membrane-bound ligands (Lemmon and Schlessinger, 2010). Ligand binding transmits conformational changes through the single transmembrane region to the cytoplasmic domain, which activates an ~300 residue tyrosine kinase module comprised of a N- and C-lobes around a central cleft which binds ATP and protein substrates (Endres et al., 2014; Maruyama, 2015; Tatulian, 2015). RTK-mediated tyrosine phosphorylation of multiple substrates activates multiple signal transduction pathways which control different cellular responses such as migration, survival, proliferation and differentiation. Although many RTK phospho-substrates have been identified with specific roles in different aspects of cellular physiology, there is no mechanism to adequately explain how RTK trafficking is regulated in resting and ligand-stimulated conditions to meter RTK bioavailability.

One RTK model is the vascular endothelial growth receptor (VEGFRs) comprising VEGFR1, VEGFR2 and VEGFR3 (Bates et al., 2018; Simons et al., 2016). VEGFR2 is a major pro-angiogenic switch which also contributes to tumor angiogenesis (Apte et al., 2019). VEGF-A binds to both VEGFR1 and VEGFR2 with different outcomes and physiological responses (Koch and Claesson-Welsh, 2012; Simons et al., 2016; Smith et al., 2015). One of the best characterized factors which interact with VEGFR2 is phospholipase Cγ1 (PLγ1), whose plasma membrane recruitment promotes PIP_2_ hydrolysis leading to cytosolic calcium ion flux and protein kinase C activation (Takahashi et al., 2001). VEGFR2 binds a number of adaptors such as TsAd (Matsumoto et al., 2005; Sun et al., 2012), Shc, Grb2, Nck (Guo et al., 1995; Kroll and Waltenberger, 1997), Crk (Stoletov et al., 2001), Shb, Sck, SHP-1, and p66Shc (Simons et al., 2016). VEGFR2 binding to epsin (Rahman et al., 2016) and synectin (Lanahan et al., 2013; Salikhova et al., 2008) suggests that interactions with endocytic regulators facilitates VEGFR2 internalization and delivery to endosomes. However, we were still lacking a mechanism to explain how VEGFR trafficking controls receptor bioavailability for exogenous VEGF-A ligand.

In this study, we explored the idea that VEGFR interaction with novel cytosolic factor(s) facilitates integration of signaling and trafficking pathways. We employed a membrane protein-based genetic screen to identify VEGFR-interacting cytosolic factors. One such protein was S100A6, a calcium-binding protein which binds both VEGFR1 and VEGFR2. Calcium-dependent S100A6 binding to VEGFR1 and VEGFR2 regulates membrane trafficking and VEGF-A-regulated signal transduction. Our model postulates a feedback circuit involving calcium-dependent protein-protein interactions which modulate VEGFR trafficking and bioavailability at the plasma membrane.

## Results

### S100A6 identified as a VEGFR2-binding protein using a membrane Y2H screen

There has been a lack of genetic screens using a native VEGFR membrane protein to identify new binding partners and potential regulators. To address this, we used the split ubiquitin membrane yeast two-hybrid (Y2H) system (Johnsson and Varshavsky, 1994; Stagljar et al., 1998) which enables the use of native membrane proteins as ‘baits’ to screen genome-wide libraries. In this membrane Y2H system, a split ubiquitin polypeptide and the LexA-VP16 transactivator were used to control nuclear yeast gene expression (*Figure 1A*). The ‘bait’ and ‘prey’ are tagged with different halves of the ubiquitin molecule, which when brought together in the cytosol, allows activation of a cytosolic ubiquitin-specific protease which cleaves the LexA-VP16 transactivator from the hybrid prey (*Figure 1A*). Cleaved LexA-VP16 can now translocate into the yeast nucleus, and stimulates auxotrophic gene expression to enable cell growth on defined media (*Figure 1A*). In this study, we fused the complete human VEGFR2 coding sequence to Cub and LexA-VP16 to form a ‘bait’ hybrid protein to screen for binding to interacting factors (protein X) fused to the NubG ‘prey’ hybrid protein (*Figure 1B*).

**Figure 1.**
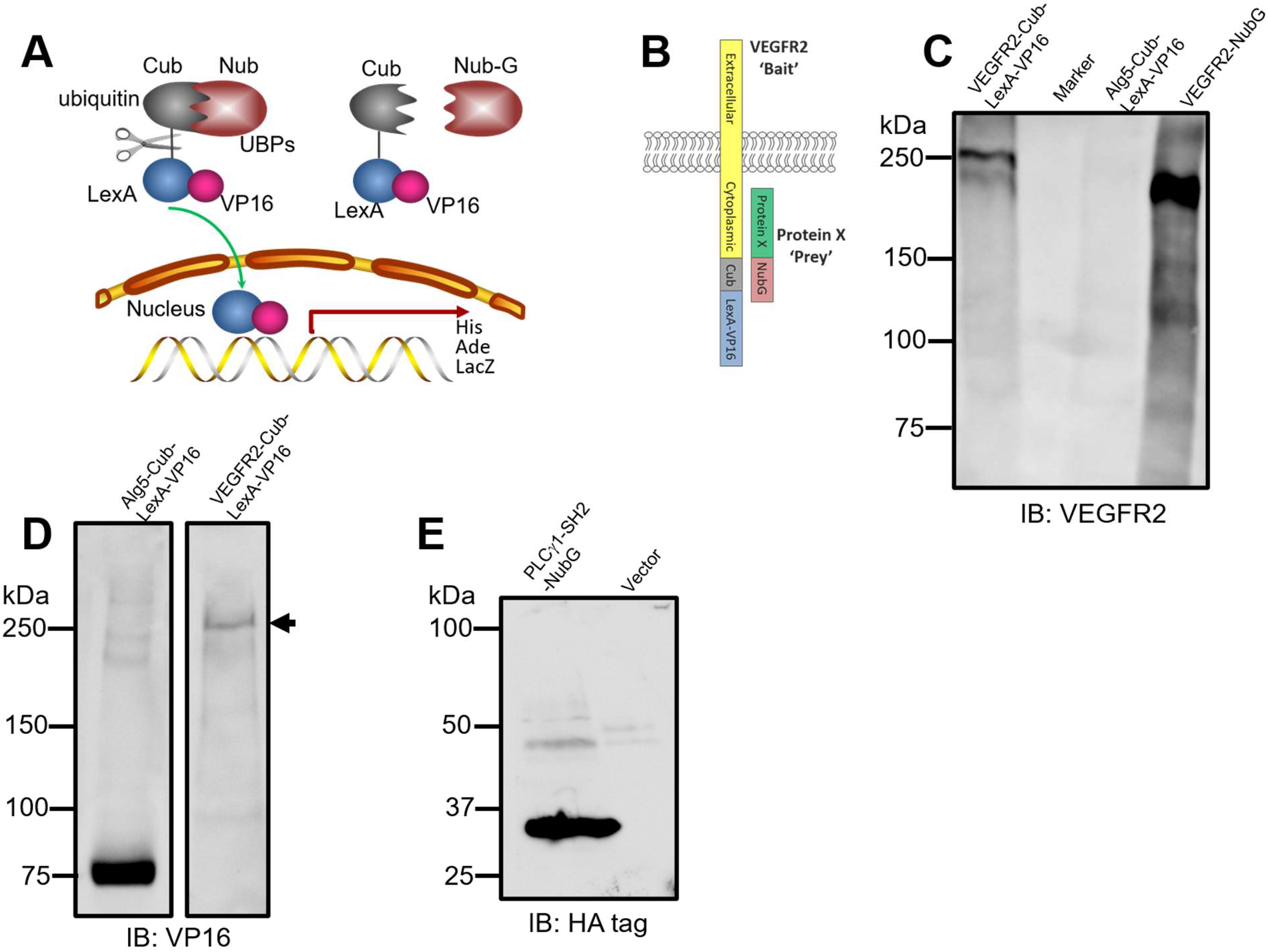
VEGFR2 expression and interaction analysis in a membrane yeast two-hybrid system. (**A**) In the split-ubiquitin system, C-terminal of ubiquitin (Cub) and N-terminal of ubiquitin (Nub) spontaneously reconstitute into a native ubiquitin fold that can be recognized by ubiquitin-specific proteases (UBPs). This UBP cleaves the artificial fusion protein to release LexA-VP16 which in turn translocates to the yeast nucleus to activate reporter gene expression. Ile>Gly substitution in Nub (Nub-G) reduces affinity for Cub, blocking spontaneous Nub-Cub re-assembly. UBP-mediated proteolysis and release of VP16 promotes transcription of auxotrophic marker genes which enable survival on histidine or adenine-deficient media. Bacterial β-galactosidase (LacZ) is an additional reporter controlled by LexA-VP16 nuclear translocation. (**B**) Schematic of the VEGFR2 bait with the full-length human VEGFR2 fused to Cub and LexA-VP16. (**C**) Expression and detection of VEGFR2-Cub-LexA-VP16 and VEGFR2-NubG (compared to control Alg5-LexA-VP16) in yeast by immunoblotting using anti-VEGFR2 antibodies. (**D**) Expression of Alg5-Cub-LexA-VP16 (control) and VEGFR2-Cub-LexA-VP16 hybrid proteins in yeast detected by immunoblotting using anti-VP16 antibodies. Arrowhead indicates full-length VEGFR2 fusion protein. (**E**) Expression of PLCγ1-SH2-NubG prey protein in yeast detected by immunoblotting using anti-HA tag antibody.

We then assessed the expression of VEGFR2 hybrid proteins using either the ‘prey’ or ‘bait’ plasmid vectors in transformed yeast cells (*Figure 1C*). Yeast expression of either VEGFR2-Cub-LexA-VP16 or VEGFR2-NubG revealed high molecular weight bands ~200-250 kDa corresponding to the predicted size of hybrid proteins (*Figure 1C*). Probing yeast cells expressing Alg5-LexA-VP16 or VEGFR2-LexA-VP16 hybrid proteins using anti-VP16 antibodies again detected bands of expected sizes (*Figure 1D*). A positive control PLCγ1-SH2 domain (known to interact with VEGFR2), with an engineered HA tag was fused to NubG, expressed in yeast cells, revealing a hybrid protein of expected size (*Figure 1E*). This PLCγ1-SH2-NubG prey construct could now be used as a positive control in subsequent yeast genetic screens.

We checked for yeast reporter gene expression comparing bait VEGFR2 with positive and negative controls (*Supplement Figure S1*). We used the PLCγ1-SH2 domain that binds the phosphotyrosine epitope in the VEGFR2 cytoplasmic tail (Guo et al., 1995; Takahashi et al., 2001); (Larose et al., 1995). When fused to the Nub-G prey construct, PLCγ1-SH2-Nub promoted a 4-5-fold increase (vs. controls) in LacZ (β-galactosidase) activity (*Supplement Figure S1*), suggesting interaction between VEGFR2 ‘bait’ and PLCγ1-SH2 ‘prey’. We also tested whether VEGFR2 could form homodimers by co-expressing a VEGFR2-NubG ‘prey’ construct. Again, there was a 5-fold increase in LacZ activity (*Supplement Figure S1*), indicating that VEGFR2-VEGFR2 homodimers were formed. When control yeast proteins such as Fur4, Ost1 and Alg5 were fused to NubG, LacZ expression was relatively low but higher than ‘empty’ prey vector (Nub-G) alone. However, when the same proteins are fused to NubI, which causes self-association with the Cub moiety on the bait protein, LacZ expression was increased 5-fold (*Supplement Figure S1*).

We constructed and screened a ‘prey’ cDNA library of human endothelial proteins fused to NubG-LexA-VP16 (see Materials and Methods). From this screen, we identified a calcium-binding protein S100A6 as a potential binding partner for VEGFR2. The S100A6 prey plasmid construct was isolated, re-transformed into yeast cells and compared to a range of controls under defined growth conditions (*Supplement Table 1*). The S100A6 prey showed yeast growth when co-expressed with VEGFR2 bait (*Supplement Table 1*), indicating protein-protein interactions between the two molecules. LacZ activity assay showed that co-expression of VEGFR2 bait and S100A6 prey caused a 5-fold rise in LacZ activity (*Supplement Table 1*), indicating protein-protein interactions between VEGFR2 and S100A6. Biochemical analysis of S100A6-NubG in yeast cells revealed a fusion protein of the expected size that contained an engineered HA tag (*Figure S2A*) and cross-reactive with anti-human S100A6 antibodies (*Figure S2B*). Probing human endothelial cells with anti-S100A6 antibodies revealed a low molecular weight band of ~10 kDa (*Figure S2C*).

### S100A6 binds to the VEGFR2 cytoplasmic domain

Endothelial cells express both VEGFR1 and VEGFR2, two closely related but distinct gene products with distinct functional roles (Shibuya, 2015; Simons et al., 2016; Smith et al., 2015). To investigate the association of endogenously expressed VEGFRs and S100A6, we immunoisolated detergent-solubilized complexes from endothelial cells and probed for different proteins including using the transferrin receptor as a control (*Figure 2A*). As expected, VEGFR2 complexes contained S100A6; surprisingly, VEGFR1 complexes also contained S100A6 (*Figure 2A*). Immunoisolation of S100A6 complexes from endothelial cells revealed the presence of both VEGFR1 and VEGFR2 (*Figure 2A*). Another membrane protein, the transferrin receptor, was absent from immunoisolated complexes of VEGFR or S100A6 (*Figure 2A*), indicating specificity in the interaction between S100A6 and VEGFRs.

**Figure 2.**
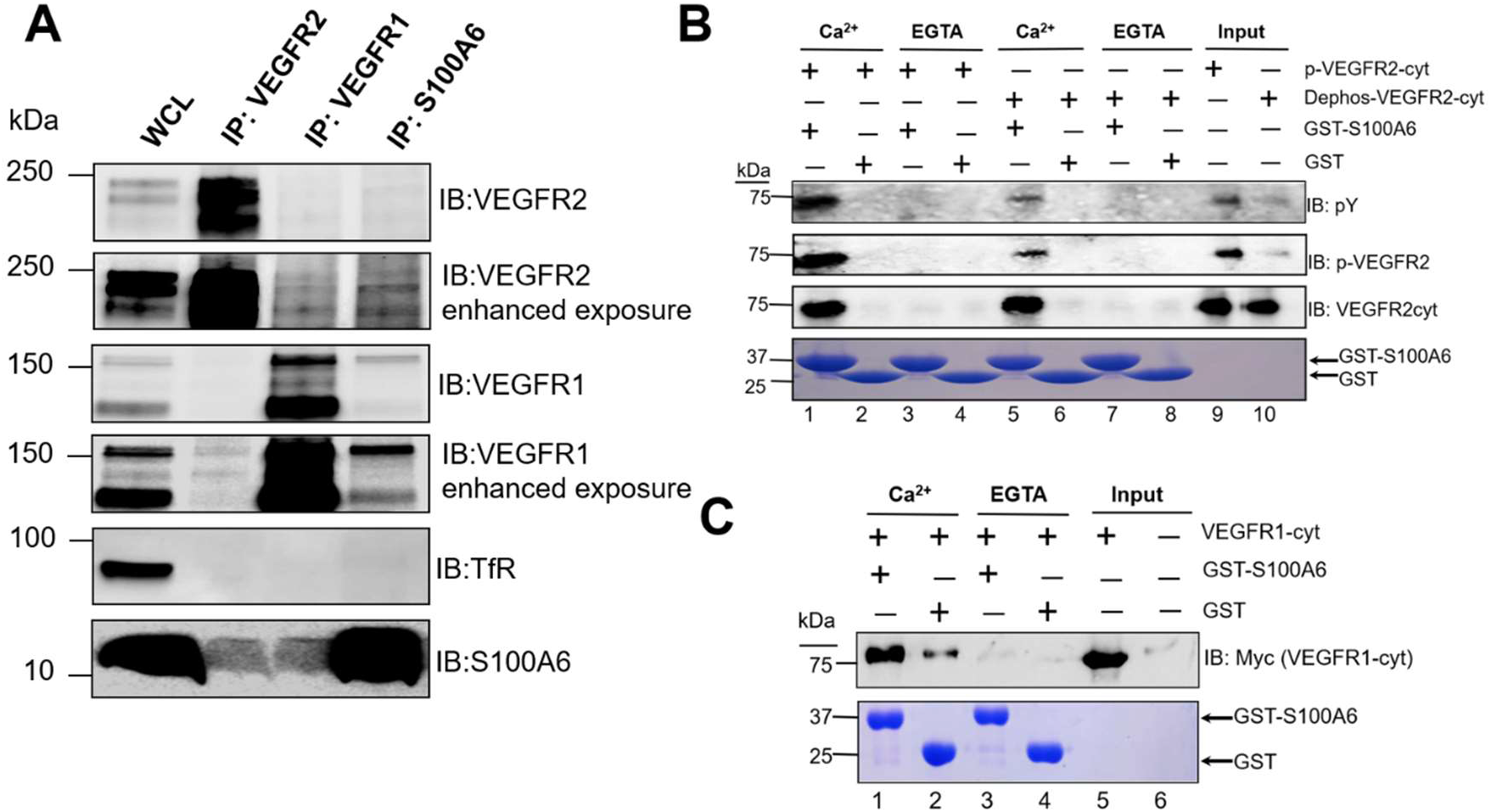
VEGFR and S100A6 complexes exhibit calcium-dependence. **(A)** Immunoisolation of VEGFR or S100A6 complexes followed by immunoblot analysis. Whole cell lysates (WCL) were subjected to detergent lysis (see Materials and Methods) and VEGFR2, VEGFR1 and S100A6 complexes isolated before SDS-PAGE and immunoblotting (IB). Goat anti-VEGFR2, goat anti-VEGFR1 or rabbit anti-S100A6 antibodies were used to isolate VEGFR or S100A6 complexes respectively. Molecular weight markers (kDa) and respective proteins are indicated in the panel. Transferrin receptor (TfR) was used as a negative control. (**B**) Recombinant proteins comprising soluble VEGFR2 cytoplasmic domain or de-phosphorylated VEGFR2 cytoplasmic domain was incubated with soluble GST-S100A6 or GST in the presence of 1 mM calcium ions or 1 mM EGTA. This was followed by incubation with glutathione-agarose beads, centrifugation and brief washes with buffer. Bound proteins were analyzed on 12% SDS-PAGE together with purified phosphorylated VEGFR2 (lane 9) or de-phosphorylated VEGFR2 (lane 10). Immunoblotting was carried out using mouse anti-phosphotyrosine (pY20), rabbit anti-VEGFR2-pY1175 or sheep anti-VEGFR2 cytoplasmic domain antibodies. (**C**) Similar experiments carried out using the soluble VEGFR1 cytoplasmic domain incubated with soluble GST-S100A6 or GST in the presence of 1 mM calcium ions or 1 mM EGTA. This was followed by incubation with glutathione-agarose beads, centrifugation and brief washes with buffer. Bound proteins were analyzed on 12% SDS-PAGE and immunoblotting using sheep anti-VEGFR1 cytoplasmic domain antibodies.

S100A6 belongs to a family of relatively small (~10 kDa) proteins which undergo calcium-dependent conformational changes which modulate protein-protein interactions (Rezvanpour and Shaw, 2009; Santamaria-Kisiel et al., 2006). One likelihood is that S100A6 interacts with the VEGFR2 cytoplasmic domain (*Figure 1B*). To test this possibility, we expressed and purified a recombinant soluble VEGFR2 cytoplasmic domain fragment to assess binding to purified recombinant S100A6 (*Figure 2B*). Two potential regulatory aspects of VEGFR2-S100A6 interactions are calcium ion binding to S100A6 (Donato et al., 2017; Santamaria-Kisiel et al., 2006) and VEGFR2 tyrosine autophosphorylation (Simons et al., 2016; Smith et al., 2015). Interestingly, recombinant VEGFR2 exhibits phosphorylation on residue Y1175, indicating functional tyrosine kinase activity (*Figure 2B*). We investigated the biochemistry of VEGFR2-S100A6 interactions: VEGFR2 cytoplasmic domain bound to S100A6 in the presence of calcium ions (*Figure 2A*). VEGFR2-S100A6 complex formation was blocked in the presence of EGTA with no evidence for VEGFR2 association with immobilized S100A6 (*Figure 2B*). We then evaluated requirement for VEGFR2 tyrosine phosphorylation in binding to S100A6: de-phosphorylated VEGFR2 still bound immobilized S100A6 in the presence of calcium ions similar to phosphorylated VEGFR2 (*Figure 2B*). Such VEGFR2-S100A6 interactions are thus calcium-dependent but do not require VEGFR2 phosphorylation.

As our data suggested that the VEGFR2 cytoplasmic domain binds to S100A6 (*Figure 1*), we asked whether the VEGFR1 cytoplasmic domain protein could also bind to S100A6 (*Figure 2B*). This VEGFR1 cytoplasmic domain protein bound to immobilized S100A6 in the presence of calcium ions (*Figure 2C*). Again, VEGFR1-S100A6 complex formation was inhibited by the addition of EGTA (*Figure 2C*). To investigate this further, we used the membrane Y2H system to assess whether VEGFR1 displayed interaction with S100A6 (*Supplement Table 1*). Yeast cells co-expressing the VEGFR1-Cub-LexA-VP16 ‘bait’ and the S100A6-Nub-G ‘prey’ showed auxotrophic growth and LacZ expression (*Supplement Table 1*). These findings are consistent with calcium-dependent VEGFR1-S100A6 complex formation.

### Biochemistry of VEGFR2-S100A6 interactions

The interaction between two molecules can be described by biochemical parameters. We explored the interactions between S100A6 and VEGFR2 using different assays (*Figure 3*). First, surface plasmon resonance (SPR) was used to measure S100A6 binding to the VEGFR2 cytoplasmic domain in the presence of calcium ions (*Figure 3A*). SPR data showed that titration of S100A6 displayed dose-dependent kinetics of binding to immobilized VEGFR2 in the presence of calcium ions (*Figure 3A*). Based on these SPR data, the VEGFR2-S100A6 dissociation constant (*K_d_*) was calculated to be ~0.2 μM. S100A6 binding to immobilized VEGFR2 was abolished in the presence of EGTA (*Figure 3A*).

**Figure 3.**
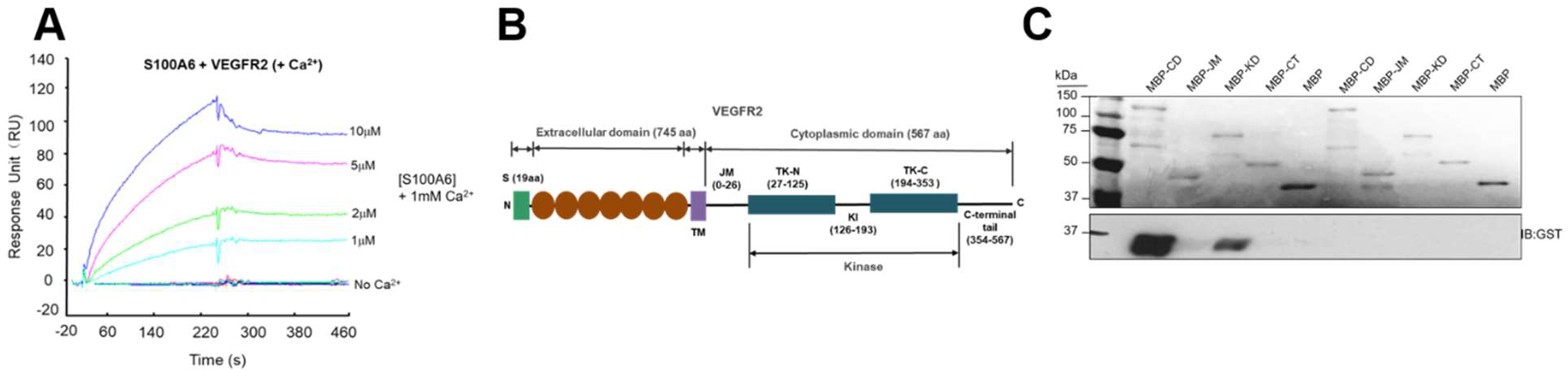
Interaction of the VEGFR2 cytoplasmic domain with S100A6. (**A**) SPR analysis of recombinant immobilized soluble VEGFR2 cytoplasmic domain binding to S100A6 the presence of 1 mM divalent calcium ions, or 1 mM EGTA (no Ca2+). Soluble VEGFR2 cytoplasmic domain was immobilized on the chip as described in Materials and Methods. GST-S100A6 solutions of 10 μM, 5 μM, 2 μM or 1 μM in buffer supplied with 1 mM calcium ions or 1 mM EGTA was flowed over immobilized VEGFR2. The response unit (RU) was recorded using evaluation software. (**B**) Schematic view of the VEGFR2 protein showing the various regions. The juxtamembrane (JM), tyrosine kinase domain (KD) and C-terminal tail (CT) are indicated on the line diagram. Sequences corresponding to residues from the VEGFR2 juxtamembrane (JM), kinase domain (KD) and C-terminal tail (CT) were fused to MBP and used in binding studies. (**C**) Interaction of MBP-VEGFR2 proteins with S100A6. The VEGFR2 cytoplasmic domain fused to MBP (Cyto), N-proximal juxtamembrane region (JM), kinase domain alone (KD) and cytoplasmic tail (CT) were tested for their ability to bind either GST or GST-S100A6 in a pull-down assay. The upper panel shows an immunoblot for GST to detect GST-S100A6 fusion; lower panel shows the Ponceau S stain showing the MBP-VEGFR2 proteins used in the assay. Molecular weight markers are indicated.

The S100A6 protein (90 residues) binding to the larger VEGFR2 cytoplasmic domain (568 residues) was mapped using deletion analysis (*Figure 3B*). We separated the 568 VEGFR2 cytoplasmic domain into the juxtamembrane (JM) region, the tyrosine kinase domain comprising the N- and C-lobes (TK-N, TK-C) including a kinase insert region, and a flexible C-terminal tail (*Figure 3B*). These different portions of the VEGFR2 cytoplasmic domain were fused to the maltose-binding protein (MBP), and MBP-VEGFR2 hybrid proteins were tested for binding to GST-S100A6 (*Figure 3C*). S100A6 protein strongly bound to the 568 residue VEGFR2 cytoplasmic domain (*Figure 3C*). Deletion analysis showed that the VEGFR2 kinase domain (329 residues) bound S100A6 (*Figure 3C*). Neither the juxtamembrane region nor the C-terminal tail showed any significant binding to S100A6 (*Figure 3C*).

### S100A6 modulates VEGFR1 and VEGFR2 trafficking, modification and turnover

VEGFR1 and VEGFR2 display complex patterns of steady-state and ligand-stimulated distribution in endothelial cells (Ewan et al., 2006; Gampel et al., 2006; Lampugnani et al., 2006; Mittar et al., 2009). One possibility is that calcium-dependent S100A6 interactions with VEGFR2 and/or VEGFR1 modulates trafficking and turnover. To assess this, we compared S100A6 with another S100 family member (S100A10) also expressed in endothelial cells (Bao et al., 2012), using protein knockdown using RNAi followed by analyses of VEGFR trafficking and cellular distribution (*Figure 4*).

**Figure 4.**
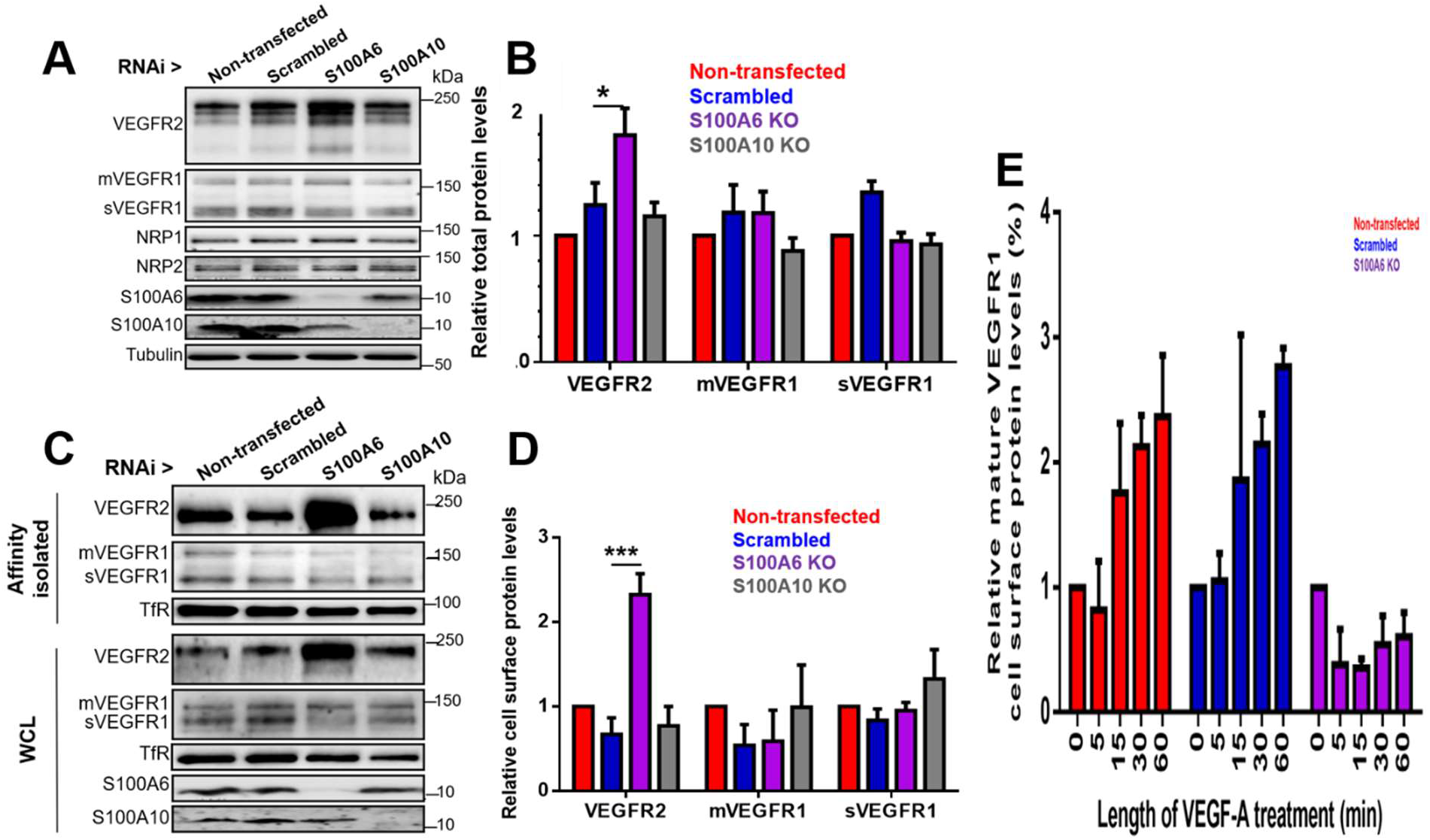
S100A6 requirement for VEGFR1 and VEGFR2 trafficking. Endothelial cells were subjected to RNAi on S100 proteins and analyzed for VEGFR2 trafficking. (**A**) Endothelial cells subjected to control, scrambled, S100A6 or S100A10 siRNA treatments were lysed and immunoblotted for various proteins indicated. Molecular weights of markers are indicated. (**B**) Quantification of relative protein levels under different conditions of RNAi. Color coding indicates non-transfected (red), scrambled siRNA (blue), S100A6 knockdown (purple) and S100A10 knockdown (grey). Error bars indicate +SEM (n>3). *, p<0.05. (**C**) Endothelial cells subjected to control, scrambled, S100A6 or S100A10 siRNA treatments followed by cell surface biotinylation, cell lysis and purification of biotinylated proteins. Whole cell lysates and purified proteins were immunoblotted for the various proteins indicated. Molecular weights of markers are also indicated. (**D**) Quantification of relative protein levels under different conditions of RNAi. Color coding indicates non-transfected (red), scrambled siRNA (blue), S100A6 knockdown (purple) and S100A10 knockdown (grey). Error bars indicate +SEM (n>3). ***, p<0.001. (E) Quantification of VEGF-A-regulated mature VEGFR1 trafficking to the plasma membrane using cell surface biotinylation. Endothelial cells subjected to control, scrambled or S100A6 siRNA treatments followed by VEGF-A (10 ng/ml) stimulation, followed by cell surface biotinylation, cell lysis and purification of biotinylated proteins before immunoblotting. Color coding indicates non-transfected (red), scrambled siRNA (blue) and S100A6 knockdown (purple).

Depletion of S100A6 caused a significant rise in overall VEGFR2 levels but this did not affect other membrane proteins including VEGFR1 or VEGF co-receptors, the neuropilins (NRP1, NRP2) (*Figure 4A*). In contrast, depletion of S100A10 did not significantly affect VEGFR or control membrane protein levels (*Figure 4A*). Quantification showed that S100A6 depletion caused ~80% rise in total VEGFR2 levels, with little or no significant effects on membrane-bound or soluble VEGFR1 levels (*Figure 4B*). We used cell surface biotinylation to assess plasma membrane VEGFR pools under these conditions (*Figure 4C*). Increased mature plasma membrane VEGFR2 levels were detected upon S100A6 knockdown (*Figure 4C*). Quantification showed ~2.5-fold increase in mature plasma membrane VEGFR2 levels compared to controls (*Figure 4D*). Under basal or resting conditions, S100A6-depleted cells showed no significant change in cell surface membrane-bound or soluble VEGFR1 compared to controls (*Figure 4D*).

A previous study showed that biosynthetic VEGFR1 undergoes VEGF-A-stimulated and calcium-dependent trafficking from the distal Golgi to the plasma membrane (Mittar et al., 2009). One possibility was that S100A6 is involved in this calcium-dependent Golgi-to-plasma membrane trafficking step. To test this idea, we used cell surface biotinylation of S100A6-depleted cells to assess VEGFR1 plasma membrane levels in resting or VEGF-A-stimulated cells (*Figure 4E*). Upon VEGF-A stimulation, we detected a time-dependent, ~2.5-fold increase in plasma membrane VEGFR1 levels over a 60 min time period (*Figure 4E*). However, S100A6 knockdown caused a complete block in VEGF-A-stimulated VEGFR trafficking to the plasma membrane (*Figure 4E*). Depletion of S100A6, but not S100A10, thus modulates both VEGFR1 and VEGFR2 trafficking and plasma membrane levels.

### S100A6 regulates VEGFR2 bioavailability and VEGF-A-stimulated signal transduction

VEGFR2 trafficking influences VEGF-A-stimulated signaling from the cell surface (Bruns et al., 2010; Ewan et al., 2006; Lampugnani et al., 2006; Manickam et al., 2011; Yamada et al., 2014). Based on our findings in this study, we then asked whether S100A6 regulation of plasma membrane VEGFR2 levels regulates VEGF-A-stimulated signal transduction events (*Figure 5*). We monitored 2 different VEGFR2 phosphotyrosine epitopes (pY1175, pY1214) which exhibit different kinetics upon VEGF-A stimulation (Fearnley et al., 2016). As expected, VEGFR2-pY1175 is rapidly generated in response to VEGF-A stimulation (*Figure 5A*). Upon S100A6 knockdown, VEGFR2-pY1175 levels are elevated (*Figure 5A*), corresponding to ~5-fold magnitude increase compared to controls (*Figure 5B*). However, the kinetics of VEGFR2-pY1175 appearance, time to peak and decline were similar in both control and S100A6-depleted endothelial cells (*Figure 5B*). Interestingly, VEGFR2-pY1214 is detected under basal conditions in line with previous studies (Fearnley et al., 2016); such signaling is also substantially elevated upon S100A6 knockdown in VEGF-A-stimulated cells (*Figure 5A*).

**Figure 5.**
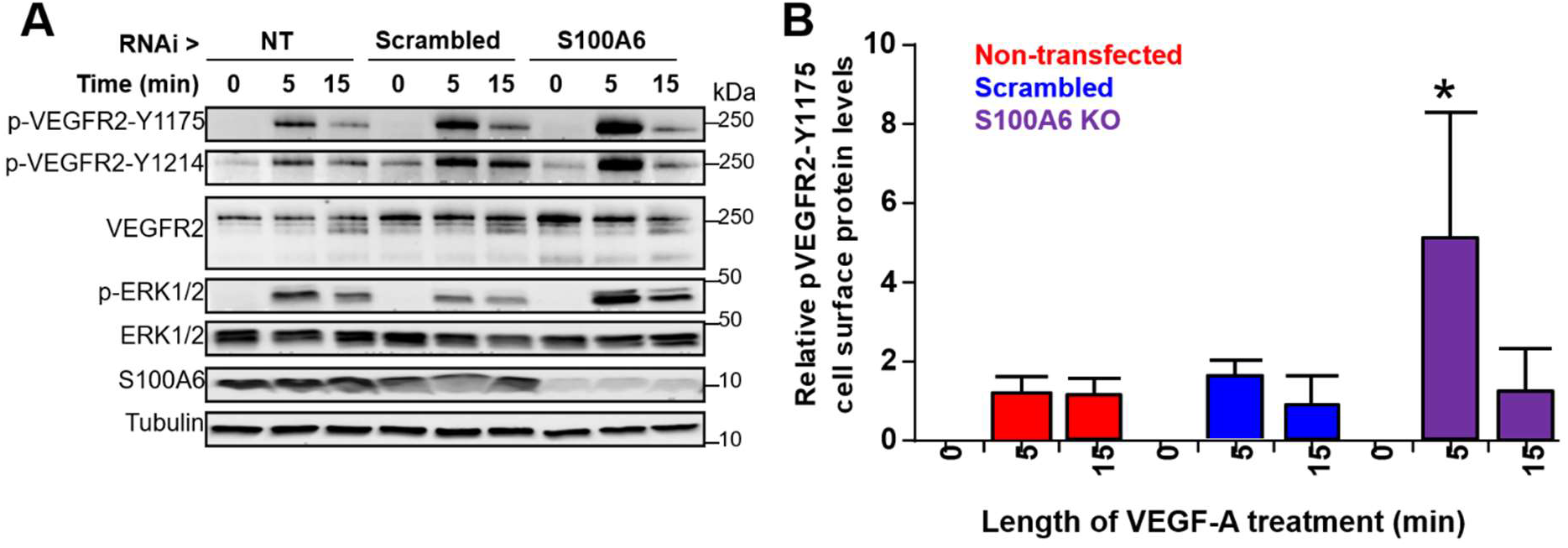
Modulation of VEGFR2 signaling by S100A6. Endothelial cells were subjected to RNA interference and analyzed by (**A**) immunoblotting, (**B**) quantification of immunoblot data. Endothelial cells subjected to control, scrambled or S100A6 siRNA treatments were stimulated with VEGF-A (10 ng/ml), lysed and immunoblotted for the VEGFR2-pY1175 epitope and relative levels quantified. The blots were also probed with anti-ERK1/2 and anti-phospho-ERK1/2 antibodies to check for canonical MAPK signaling. Blotting for tubulin was used as an additional loading control in these experiments. Error bars indicate +SEM (n>3). *, p<0.05.

We also carried out confocal microscopy to ascertain subcellular VEGFR and S100A6 localization (*Figure 6A, 6B*). Steady-state VEGFR2 distribution shows localization to the plasma membrane, endosomes and juxtanuclear Golgi region (*Figure 6A*). In contrast, S100A6 is widely distributed in the cytosol and nucleus, with occasional staining of tubular profiles emanating from the Golgi region (*Figure 6A*, short arrows). Overlay images suggests co-distribution and close proximity of VEGFR2 and S100A6 (*Figure 6A*, boxed). Analysis of the VEGFR1 and S100A6 (Figure 6B) showed also showed co-distribution in the Golgi region (*Figure 6B*, long arrows). Co-labeling of VEGFR1 and S100A6 elongated tubular profiles emanating from the Golgi region was also detected (*Figure 6B*, boxed).

**Figure 6.**
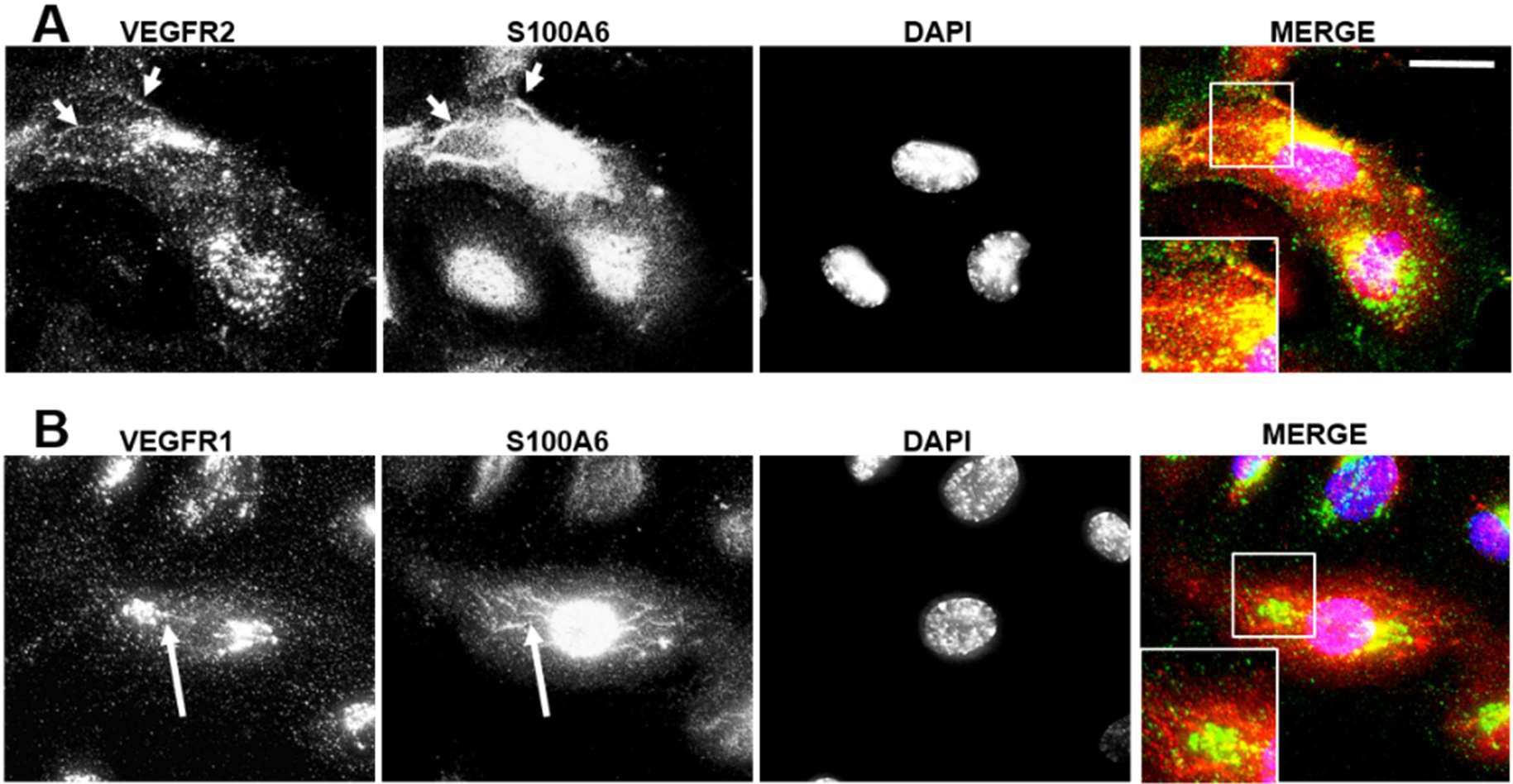
Co-distribution of VEGFRs and S100A6. Endothelial cells were fixed and processed for wide-field deconvolution microscopy analysis (see Materials and Methods) of (**A**) VEGFR2, or (**B**) VEGFR1 vs. S100A6 on fixed endothelial cells. Triple color overlay shown in extreme right-hand panel. Bar, 10 μm. Arrows and arrowheads denote co-distribution of VEGFR and S100A6 along tubular profiles. Each image shown comprises 20-35 optical sections to better visualize tubular profiles.

We then evaluated the 3-D structures of the S100A6 and VEGFR2 cytoplasmic domain using *in silico* modelling (*Figure 7*). Comparison of the structures of free and calcium-bound S100A6 shows relatively large movements of helix H3 (*Figure 7A*). *In silico* docking studies using the VEGFR2 tyrosine kinase domain suggests that calcium-bound S100A6 binds close to the cleft of the tyrosine kinase module (*Figure 7B*). It is unclear whether VEGFR tyrosine kinase activity and calcium-S100A6 binding are functionally coupled.

**Figure 7.**
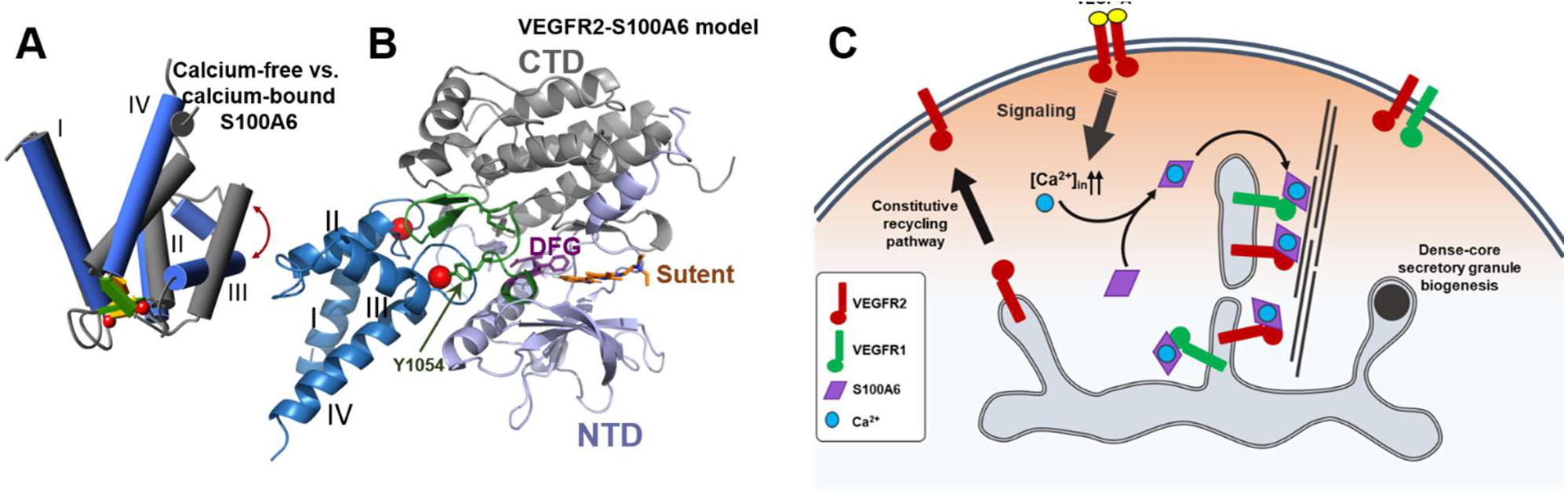
Models of VEGFR2/S100A6 interaction and membrane trafficking. (**A**) Crystal structures of calcium-free (grey: PDB ID 1K9P) and calcium-bound (blue: PDB ID 1K96) forms of S100A6. There is a large movement of Helix III (brown arrow) upon binding Ca2+ (red spheres). (**B**) Energy minimized model of VEGFR2 kinase domain (PDB ID: 4AGD) bound to calcium-bound S100A6 (PDB ID: 1K96) using in silico docking with HADDOCK 2.2 (see Materials and Methods). The kinase domain insert between the N-terminal (NTD: light blue) and C-terminal (CTD: grey) segments of the kinase domain is not resolved in VEGFR2 tyrosine kinase domain structure (PDB ID: 4AGD). The regulatory loop that contains the autophosphorylation sites Y1054 and Y1059 is shown green, with the conserved DFG motif highlighted (purple). Sunitinib (Sutent: orange) is bound to the kinase domain. The major contacts contributed by S100A6 are via the interhelical Ca2+-binding loops. (**C**) Regulation of TGN-to-plasma membrane trafficking of VEGFR1 and VEGFR2 requiring S100A6. Depicted are 3 parallel transport routes: a constitutive transport step accessed by VEGFR2, a calcium-dependent trafficking route from the TGN which is dependent on S100A6 binding to the cytoplasmic domains of VEGFR1 and VEGFR2 cargo for inclusion into a new class of transport carriers, and the dense-core secretory granule route.

## Discussion

How does a cell integrate membrane receptor bioavailability for a specific ligand? In the case of VEGF-A binding to two different membrane receptors, VEGFR1 and VEGFR2, different pathways of signaling, trafficking and turnover need to be integrated to control cellular responses such as cell migration, proliferation and tubulogenesis. Up to now, we lacked molecules that could bridge VEGFR signaling and trafficking. Herein, we now present evidence that a calcium-dependent cytosolic protein, S100A6, binds both VEGFR1 and VEGFR2 to integrate signaling and trafficking pathways. Five lines of evidence support this conclusion. Firstly, a genetic screen of human endothelial proteins identified S100A6 as a binding partner for VEGFR2. Second, membrane Y2H assay shows that either VEGFR2 or VEGFR1 can interact with S100A6. This was confirmed by the detection of stable complexes of S100A6 with either VEGFR1 or VEGFR2 in endothelial cells. Third, S100A6 binds the VEGFR2 cytoplasmic domain *in vitro*, with sub-micromolar binding affinity (*K_d_*) and displays calcium-dependence. Furthermore, the VEGFR1 cytoplasmic domain also binds to S100A6 in a calcium-dependent manner. S100A6 binding to VEGFR2 maps to the tyrosine kinase module. Fourth, S100A6 regulates VEGFR1 and VEGFR2 trafficking with different functional outcomes. Whereas S100A6 depletion causes dysregulated VEGFR2 trafficking and increased plasma membrane levels, loss of S100A6 completely blocks VEGFR1 Golgi-to-plasma membrane trafficking. Finally, S100A6 influences VEGFR2 plasma membrane bioavailability by modulating VEGF-A-regulated VEGFR2 tyrosine phosphorylation and downstream canonical MAPK signaling.

Our study supports a mechanism where at least two trafficking routes regulate VEGFR delivery from the *trans-*Golgi network (TGN) to the plasma membrane (*Figure 7C*). In higher eukaryotes, a constitutive TGN-to-plasma membrane anterograde trafficking step is utilized by a majority of soluble and membrane-bound secretory proteins (Guo et al., 2014; Pakdel and von Blume, 2018). Our findings that S100A6 depletion leading to elevated VEGFR2 levels at the plasma membrane suggests dysregulation in TGN-to-plasma membrane trafficking. One explanation is that VEGFR2 utilizes both constitutive and calcium-regulated trafficking routes to exit the TGN (Figure 7C). Newly synthesized VEGFR2 also accumulates within the Golgi (Manickam et al., 2011); it was postulated constitutive TGN-to-plasma membrane trafficking enables replenishment of the cell surface VEGFR2 pool undergoing endocytosis, recycling or degradation (Ewan et al., 2006; Jopling et al., 2011). Our studies now suggest that plasma membrane VEGFR2 bioavailability is also dependent on calcium-regulated TGN-to-plasma membrane trafficking event (*Figure 7C*). VEGFR2 levels are thus ‘metered’ by 2 parallel anterograde trafficking steps to ensure that biosynthetic VEGFR2 delivery is synchronized with endocytosis of plasma membrane VEGFR2 for delivery to endosomes.

Endothelial cells display VEGFR2 localization to Golgi, plasma membrane and endosomes (Ewan et al., 2006; Gampel et al., 2006; Manickam et al., 2011). Co-distribution of VEGFR1, VEGFR2 and S100A6 on tubular profiles emanating from a juxtanuclear Golgi region in endothelial cells indicate transport carriers which mediate VEGFR cargo delivery to the plasma membrane. VEGFR2 Golgi trafficking shows dependence on cytosolic factors such as t-SNARE syntaxin 6 (Manickam et al., 2011), the KIF13B microtubule motor (Yamada et al., 2014) and the Myo1c actomyosin motor (Tiwari et al., 2013). However, the cytoskeletal machinery involved in this calcium-regulated TGN-to-plasma membrane trafficking step (*Figure 7C*) is at present ill-defined.

VEGFR1 intracellular localization is complicated by a lack of clear functional roles for this RTK. Generally, VEGFR1 is thought to act as a ‘VEGF sink’ which sequesters ligand and acts as a negative regulator of angiogenesis, but also modulates some aspects of cancer cell proliferation (Autiero et al., 2003; Jones et al., 2009; Lichtenberger et al., 2010; Yang et al., 2006). Different studies suggest VEGFR1 localization to Golgi (Mittar et al., 2009) and nuclear (Lee et al., 2007; Zhang et al., 2010) compartments. However, VEGFR1 plasma membrane levels are elevated by cytosolic calcium ion flux in endothelial cells (Mittar et al., 2009) and cardiomyocytes (Yang et al., 2015), suggesting that VEGFR1 can undergo calcium-dependent TGN-to-plasma membrane trafficking in different cell types. Our study now provides a mechanism to explain this phenomenon with the finding that calcium-dependent binding of S100A6 to the VEGFR1 cytoplasmic domain. Depletion of S100A6 levels blocked VEGF-A-stimulated VEGFR1 delivery to the plasma membrane, consistent with VEGFR1 cargo sequestration into calcium-regulated TGN-to-plasma membrane transport carriers (*Figure 7C*). Interestingly, the VEGFR1 Golgi pool shows partial co-distribution with TGN46, a standard marker for the human TGN (Mittar et al., 2009). One explanation is that the mammalian TGN is more extensive than currently postulated, with steady-state VEGFR1 residence within a TGN-like subcompartment. From this location VEGFR1 membrane cargo is delivered to the plasma membrane via this S100A6-regulated trafficking step.

How can our proposed model (*Figure 7C*) be reconciled with other TGN trafficking events? The TGN is a site of multiple sorting, packaging and transport events, including constitutive and regulated secretion (Guo et al., 2014). In endothelial cells, the biogenesis of electron dense cylindrical Weibel-Palade bodies (WBPs) from the TGN precedes requirement for a calcium-regulated stimulus to undergo docking and fusion with the plasma membrane (McCormack et al., 2017). However, there is little or no evidence of VEGFR1 or VEGFR2 association with WBP trafficking. This then raises the question whether such a calcium-regulated trafficking event (*Figure 7C*) is endothelial-specific or exists in other cell types. Interestingly, there are similarities to calcium-regulated TGN dynamics in immortalized cell lines (Pakdel and von Blume, 2018; von Blume et al., 2012) and neurons (Mikhaylova et al., 2010; Mundhenk et al., 2019). The TGN resident and calcium-binding protein Cab45 is in close proximity to a calcium pump, SPCA1 (Deng et al., 2018; von Blume et al., 2012), but how this is linked to constitutive or regulated protein secretion was unknown. Recent studies suggest another member of the S100 family, S100A10, is involved in Weibel-Palade body exocytosis (Chehab et al., 2017). Our studies now suggest existence of a specialized calcium-regulated trafficking route from the TGN-to-plasma membrane in higher eukaryotes.

S100A6-VEGFR interactions involves binding to the tyrosine kinase region (*Figure 7B*). Deletion analysis of the VEGFR2 cytoplasmic domain maps calcium-S100A6 binding to the tyrosine kinase module. In this context, de-phosphorylation of the VEGFR2 cytoplasmic domain at Y1175 (within the carboxy-proximal tail region) does not significantly affect VEGFR2-S100A6 complex formation. Interestingly, VEGFR2 undergoes tyrosine phosphorylation at 6-8 distinct epitopes, some of which are present within the tyrosine kinase domain. It remains to be determined whether other phosphotyrosines hinder or promote S100A6 recruitment, and thus influence VEGFR TGN-to-plasma membrane trafficking. Recently, it was reported that SUMOylation of the VEGFR2 cytoplasmic domain mediates Golgi targeting (Zhou et al., 2018). SUMOylation of VEGFR2-K1270 within the flexible carboxy-terminal tail (Zhou et al., 2018) is unlikely to modulate S100A6 binding to the tyrosine kinase region (residues 833-1162; *Figure 7B*) in this context.

Our study provides a new mechanism where newly synthesized membrane cargo trafficking to the plasma membrane is dependent on integration with signal transduction pathways. Here, activation of plasma membrane receptors which trigger rise in second messenger levels such as calcium ions causes conformational changes in S100A6 to enable binding to VEGFRs, and cargo selection for TGN-to-plasma membrane trafficking. Importantly, our findings also provide a mechanistic explanation for how trafficking and secretion of newly synthesized membrane cargo is synergized with plasma membrane signaling for replenishment of membrane receptors. In this context, there is increasing evidence that S100 protein family members e.g. S100A10, regulates biosynthetic trafficking of sodium (Okuse et al., 2002) and potassium (Girard et al., 2002) channels, and regulates Weibel-Palade body exocytosis (Chehab et al., 2017).

Our proposed mechanism enables the endothelial cell to integrate plasma membrane VEGFR2 activation, downstream signal transduction with secretion of newly synthesized VEGFRs to regulate plasma membrane VEGFR bioavailability for VEGF-A. This calcium-regulated mechanism acts as a feedback loop that synchronizes VEGFR2 activation with controlling arrival of both VEGFRs at the cell surface, thus enabling tight control of cellular responses to exogenous VEGF-A. One of the puzzles in understanding VEGF biology is the existence of 2 receptor tyrosine kinases (VEGFR1, VEGFR2) that bind with differing affinity to the same ligand, VEGF-A (Ewan et al., 2006; Vaisman et al., 1990). VEGFR2 plays a major role in angiogenesis but the role of VEGFR1 is less clear (Shibuya and Claesson-Welsh, 2006). VEGFR1 is postulated to negatively regulate angiogenesis by acting as a ‘VEGF trap’; however, VEGFR1-specific ligands such as PlGF are functionally implicated in some cancers (Fischer et al., 2008). One likelihood is that calcium-stimulated VEGFR1 arrival at the cell surface (and higher affinity for VEGF-A) dictates preferential sequestration of VEGF-A, thus reducing VEGF-A bioavailability to VEGFR2. VEGFR1 could thus not only act as a ligand trap under these situations, but have a different signaling role distinct from the pro-angiogenic VEGFR2. In tumor angiogenesis, pathological levels of exogenous VEGF-A could not only promote calcium-stimulated translocation of both VEGFR1 and VEGFR2 to the plasma membrane, but continue to promote pro-angiogenic signaling through VEGFR2. Future work is needed to explore this mechanism in the context of health and disease states.

## Materials and Methods

### Gene Manipulation

The full-length human VEGFR2 open reading frame was amplified using PCR using pVEGFR2-EGFP (Jopling et al., 2011). The PLCγ1-N-SH2 domain (Larose et al., 1995) was amplified by using pGEX-2T-PLCγ1-N-SH2 plasmid. Full-length VEGFR2 was cloned into both the pBT3-SUC ‘bait’ and pPR3-N ‘prey’ plasmids. The PLCγ1-SH2 domain was cloned into pPR3-N ‘prey’ plasmid. Control Fur4-NubG, Ost1-NubG, Fur4-NubI, Ost1-NubI, pBT3-SUC2, pPR3-N, Alg5-NubG, Alg-NubI plasmids were already provided as controls. All recombinant constructs were checked by DNA sequence analysis. Total endothelial RNA was extracted from confluent early passage endothelial cells using Trizol (Sigma-Aldrich). 1^st^ strand cDNA was synthesized using SMART (Switching Mechanism At 5’ end of the RNA Transcript) technique using a kit (Easyclone cDNA Synthesis). The ds cDNA fragments with SfiI restriction enzyme sites was directionally cloned into the pPR3-N ‘prey’ plasmid which will express NubG-cDNA ‘prey’ fusion proteins. This cDNA library was transformed into yeast strain NMY51 containing the VEGFR2 bait by the lithium acetate transformation method. The colonies grown on media lacking histidine and adenine (SD-LWHA) plate were tested for β-galactosidase activity using an X-gal filter test and further β-galactosidase activity assay. Plasmids isolated from yeast colonies expressing β-galactosidase were transformed into *E. coli* for plasmid purification and DNA sequencing. DNA sequence analysis was carried out using software packages at the European Bioinformatics Institute (Hinxton, UK).

### Construction of Endothelial cDNA Library and Membrane Y2H analysis

Total RNA was extracted from confluent early passage (P0-P2) endothelial cells using Trizol (Sigma-Aldrich). 1^st^ strand cDNA was synthesized using the Easyclone cDNA library kit with SMART (Switching Mechanism At 5’ end of the RNA Transcript) technique. The cDNA fragments with SfiI restriction enzyme sites were directionally cloned into the pPR3-N ‘prey’ plasmid which expressed these NubG-cDNA constructs as ‘prey’ hybrid proteins. The cDNA library were transformed into NMY51 *S.cerevisiae* strain containing VEGFR2 bait construct by the lithium acetate transformation. Yeast colonies which grew on media lacking histidine and adenine (SD-LWHA) were picked and re-checked for β-galactosidase (LacZ) expression using the X-gal filter test. Plasmids were isolated from blue X-gal positive colonies and retransformed into the XL-1 Blue *E. coli* strain. Positive plasmids were sequenced and re-checked by rounds of transformation, expression and verification in the yeast assay. Quantification of β-galactosidase (LacZ) expression was done by picking several yeast colonies for each interaction pair. A high throughput β-galactosidase activity kit was used to evaluate expression of this marker.

### Pull-down assays for protein-protein interactions

Bacterial cultures of 50-500 ml expressing GST, MBP or hexahistidine-tagged constructs were lysed in buffer (1% (w/v) Triton X-100, protease cocktail inhibitors,1 mM PMSF in PBS), then subjected to rapid purification by sonication before incubation with glutathione-agarose, amylose resin or nickel-agarose beads at 4°C for 2 h. Beads were washed 3 times, and purified proteins eluted using glutathione, maltose or imidazole respectively. In binding studies, immobilized proteins were not eluted but incubated with purified test protein in non-ionic detergent buffer (150 mM NaCl, 10 mM Tris, 1% (w/v) NP-40, 1 mM Na_3_VO_4_, 10 mM NaF, 1 mM CaCl_2_ or 1 mM EGTA, protease cocktail inhibitor mix, 1 mM PMSF) for 2 h at 4°C. These were then briefly centrifuged and washed with lysis buffer 3 times. 2X Sample buffer containing 5% β-mercaptoethanol was added, followed by boiling and SDS-PAGE. Gels were either subjected to immunoblotting using appropriate antibodies or stained using R-250 Coomassie brilliant blue.

### Surface plasmon resonance (SPR)

Surface plasmon resonance was used to analyze protein-protein affinity. EDC (N-ethyl-N’-(dimethylaminopropyl)-carbodiimide) / NHS (N-hydroxy- succinimide) was used to activate a CM5 sensor chip flow cells #1-3. Purified soluble VEGFR2 (0.2 mg/ml in 200 mM sodium acetate, pH 5.5) was injected onto flow cell 2 to generate surface densities of 4000 RU. Rabbit anti-GST antibody in 0.2 M sodium acetate buffer was injected onto flow cell #3 to generate surface densities of 19 000 RU. Flow cell 1 was used as an activated blank flow cell. Subsequently 1 M ethanolamine pH 8.5 was injected onto each flow cell to quench unreacted esters. Different concentrations of recombinant GST-S100A6 or GST in running buffer (10 mM Tris-HCl pH 7.3, 150 mM NaCl, 0.005% (v/v) surfactant P20) containing 1 mM EDTA or 1 mM CaCl_2_ was injected for 5 min at a flow rate of 10 μl/min sequentially over the three flow cells. Association and dissociation phases were recorded for 5 min for each reaction. Every cycle was finished with two injections of 10 mM glycine pH 1.9 (20 sec) to remove non-covalently bound proteins from the chip surface. Non-specific binding was subtracted from the activated blank flow cell for every cycle. The analyses were performed on a Biacore 3000 system and the data were evaluated with BIAevaluation 3.2 software.

### Cell culture, protein knockdown and immunoblotting

Human umbilical vein endothelial cells (HUVECs) were cultured and analyzed as previously described (Fearnley et al., 2014). For immunoprecipitation of protein complexes from HUVECs were lysed in buffer containing 1% (w/v) n-dodecyl-β-D-maltoside, 150 mM NaCl, 25 mM Tris pH 7.5, 1 mM Na_3_VO_4_, 10 mM NaF, 1 mM CaCl_2_, protease cocktail inhibitors and 1 mM PMSF. 0.5-1 μg of purified goat anti-VEGFR2, goat anti-VEGFR1 or mouse anti-S100A6 antibodies were added to the soluble supernatant. After 16-24 h incubation, protein G-agarose beads were added for another 2 h, briefly centrifuged and washed 3 times with lysis buffer. Samples were resuspended in 2X reducing sample buffer, boiled and subjected to SDS-PAGE and immunobotting. For signaling experiments, HUVECs were starved for 2 h in media lacking growth factors but containing 0.2% (w/v) BSA. VEGF-A (10 ng/ml) was added for different times and cells lysed in 2% SDS, PBS, protease cocktail inhibitors, 1 mM PMSF. 20 μg of total cell lysate was subjected to SDS-PAGE and immunoblotting. Band intensities were quantified using a digital system as previously described(Fearnley et al., 2014). For cells subjected to RNAi, endothelial cells were incubated under mock-transfection conditions or incubated with 10 nM siRNA duplex as previously described (Fearnley et al., 2016; Smith et al., 2017). Cell surface biotinylation and analysis was carried out as previously described (Fearnley et al., 2016).

### Confocal microscopy

HUVECs were grown on glass coverslips and fixed with 3% (w/v) paraformaldehyde, quenched with 50 mM ammonium chloride, washed with PBS and subjected to a 5 min permeabilization with 0.2% (w/v) TX-100. Non-specific binding sites were blocked by incubation with 0.2% BSA/PBS blocking buffer before incubation in primary antibodies such as goat anti-VEGFR1 or goat anti-VEGFR2 in combination with mouse anti-S100A6 (1 @g/ml) for 16-20 h. After extensive washes, cells were incubated with AlexaFluor 488 donkey anti-goat and AlexaFluor 594 donkey anti-mouse conjugates (1 @g/ml) for 1-2 h. After washes, coverslips were inverted on a drop of mounting medium and sealed on a glass slide. Samples were viewed using a DeltaVision wide-field deconvolution microscope and 0.365 μm optical sections collected as previously described (Mittar et al., 2009). Each wide-field image shown is comprises a stack of 20-35 optical sections to better visualize 3-D structures such as tubules.

### S100A6 and VEGFR2 modeling

A structural model of the VEGFR2 tyrosine kinase domain (PDB ID:1VR2) (McTigue et al., 1999) or bound to the tyrosine kinase inhibitor Sunitinib (PDB ID: 4AGD) was used to dock the human calcium-bound S100A6 (PDB ID: 1K96) and calcium-free (PDB ID: 1K9P) (Otterbein et al., 2002). Protein structures were prepared for docking using Biovia Discovery Studio Modelling Environment, Release 3.5 (Biovia Software Inc., San Diego). A model for the complex was developed using the high ambiguity-driven protein-protein docking approach, HADDOCK (Dominguez et al., 2003), for *in silico* docking of the individual protein structures (using the experimentally determined structures). The Naccess program (Hubbard and Thornton, 1993) was utilized to analyze the solvent-accessible residues which were defined as the active residues for the docking protocol. Default Haddock parameters were used. The resultant docked poses were analyzed using Biovia Discovery Studio 3.5 for inter-molecular interactions. Further modelling studies were carried out using HADDOCK 2.2 (van Zundert et al., 2016) and CPORT (de Vries and Bonvin, 2011).

### Quantification and statistical analysis

We used one-way analysis of variance (ANOVA) followed by Tukey’s post-hoc test or two-way ANOVA followed by Bonferroni multiple comparison test using GraphPad Prism software (La Jolla, USA). Significant differences between control and test groups were evaluated with *P* values less than 0.05 (*), 0.01 (**), 0.001 (***) and 0.0001 (****) indicated on the graphs. Error bars in histograms denote mean ±SEM.

### Data availability

The study did not generate large datasets. All necessary data is included in the figures in this study.

## Abbreviations

VEGF-A: Vascular endothelial growth factor A
VEGFR: Vascular endothelial growth factor receptor
S100A6: Calcium-binding S100 protein A6

## Materials

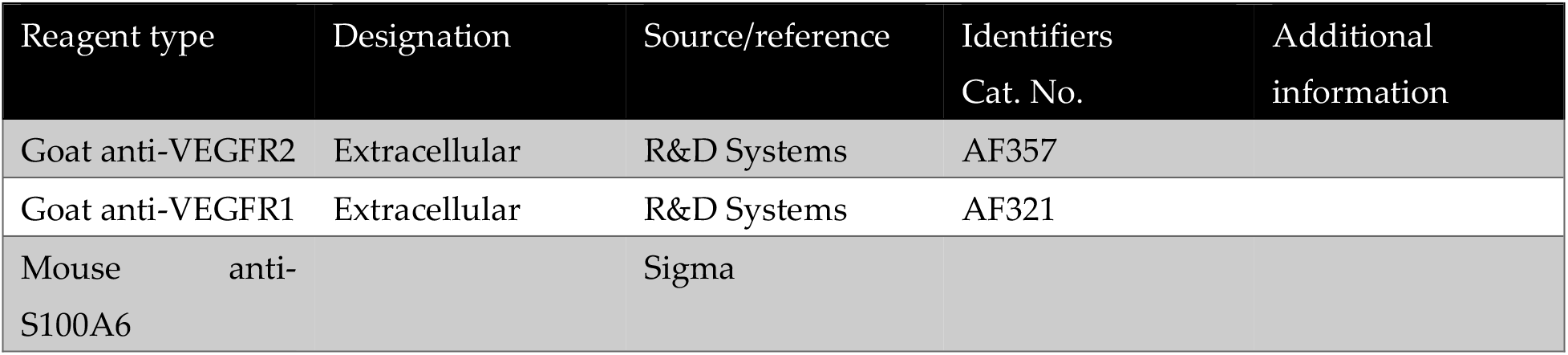
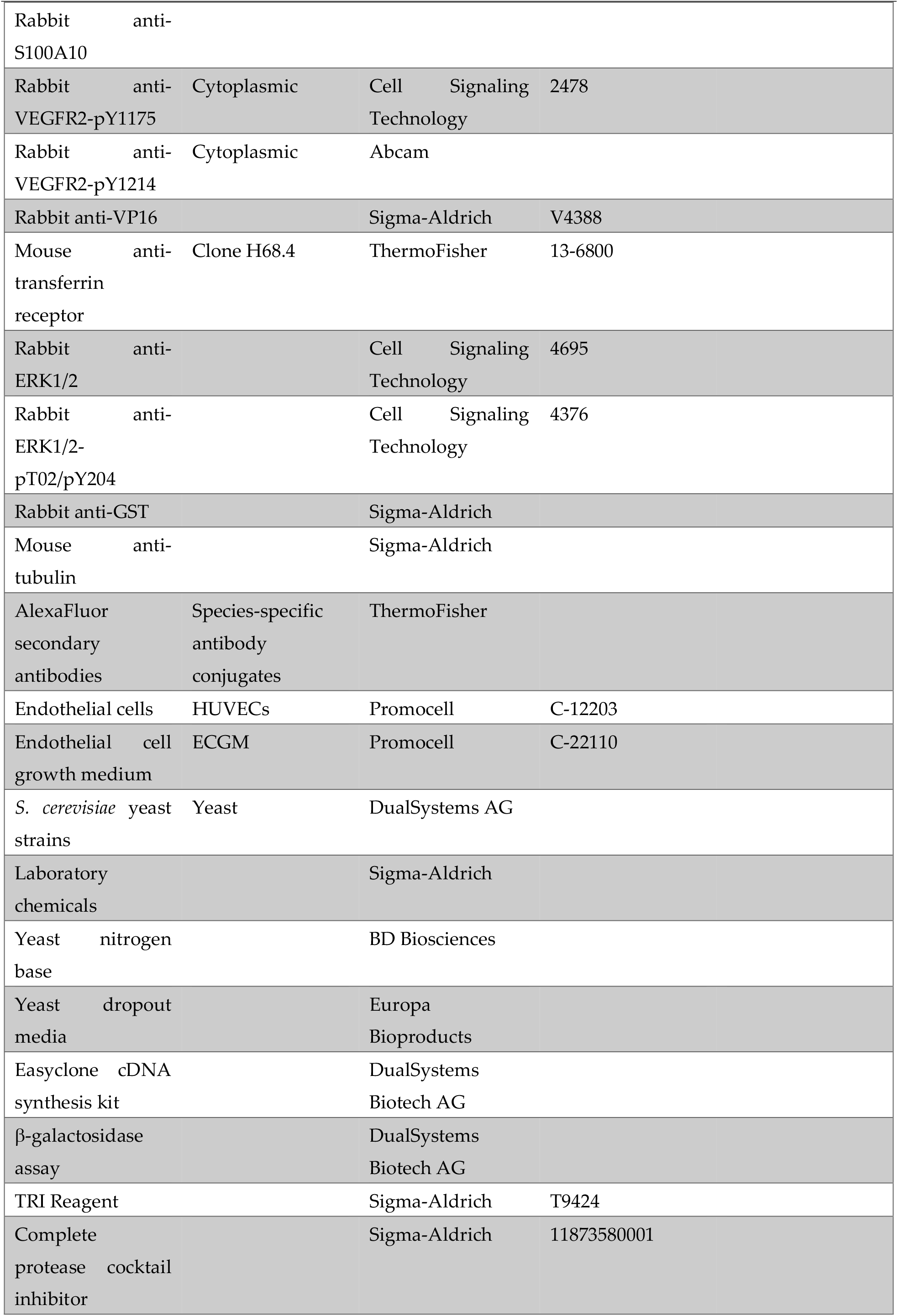
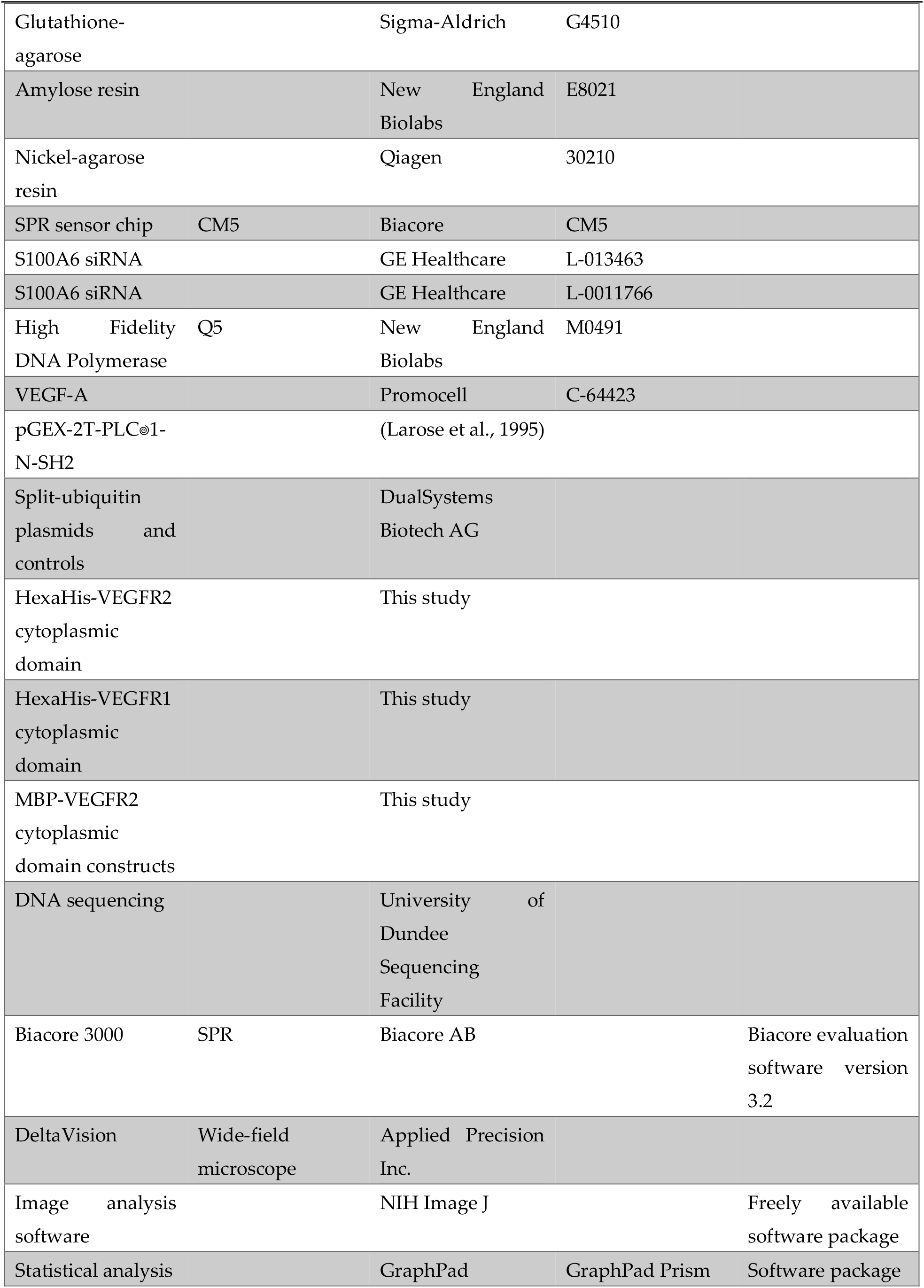

### Supplementary Materials

The following supplementary data and tables are available online at XXX.

## Author contributions

LB, GWF, CCL, AFO, acquisition of data, conception and design, analysis and interpretation of data; ACR, JBCF, contributed essential data and reagents, data analysis and interpretation, drafting and revising the manuscript; GKK, MAH, design and analysis of structural biology and modeling; ACR, GKK, MAH, JEL, SP, drafting and revising the article.

## Acknowledgments

We thank Iain Manfield (Protein Analysis Facility) for help with the SPR analysis. We thank members of the Endothelial Cell Biology Unit for help, advice and comments on the manuscript. We dedicate this manuscript to our retired colleague, John Walker, for his advice and insights into calcium-binding proteins, his unfailing good cheer and support.

## SUPPLEMENTARY MATERIALS

**Table 1.** Membrane Y2H analysis of VEGFR2 and VEGFR1 interaction with S100A6. Abbreviations for yeast minimal (SD) culture medium additives: L, Leucine; H, Histidine; A, Adenine; W, Tryptophan. All bait plasmids were co-transformed with test prey plasmids, either as empty vector carrying only NubG (pR3-N) or carrying ORFs for denoted NubG or NubI hybrid proteins. Fur4 yeast plasma membrane protein expression was used as another negative control. +++/++/+ denotes scoring of colony growth 5 days post-transformation.

| Bait | Prey | -LW | -LWH | -LWHA |
| --- | --- | --- | --- | --- |
| VEGFR2-Cub | pPR3-N (vector only) | +++ | - | - |
| VEGFR2-Cub | Alg5-NubI | +++ | ++ | + |
| VEGFR2-Cub | Fur4-NubG | +++ | - | - |
| VEGFR2-Cub | PLCg1-(SH2)-NubG | +++ | + | - |
| VEGFR2-Cub | S100A6-NubG | +++ | + | - |
| VEGFR1-Cub | pPR3-N (vector only) | +++ | - | - |
| VEGFR1-Cub | Alg5-NubI | +++ | ++ | + |
| VEGFR1-Cub | Fur4-NubG | +++ | - | - |
| VEGFR1-Cub | S100A6-NubG | +++ | + | - |
| pMBV-Fur4 | S100A6-NubG | +++ | - | - |

**Figure S1.**
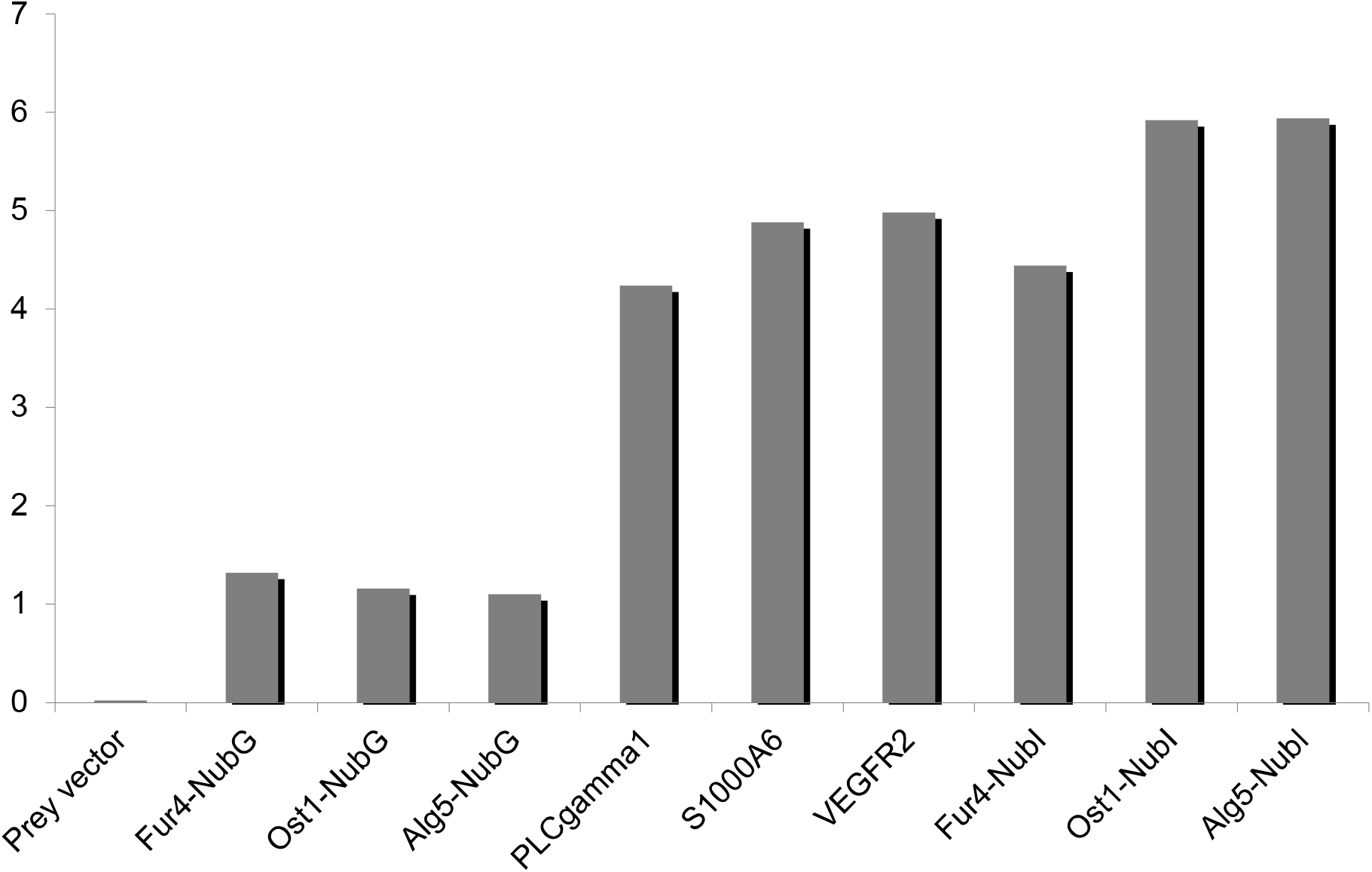
Measurement of β-galactosidase reporter activity with different bait and prey combinations. Co-expression of VEGFR2 bait with control NubG (alone) or NubG fused to Alg5, Ost1 or Fur4 (negative controls) or NubI fused to Alg5, Ost1 or Fur4 (non-specific control). Co-expression of VEGFR2-Cub bait with VEGFR2, PLCγ1-SH2 or S100A6 test proteins fused to NubG. LacZ activity assay representative of 3 or more independent experiments.

**Figure S2.**
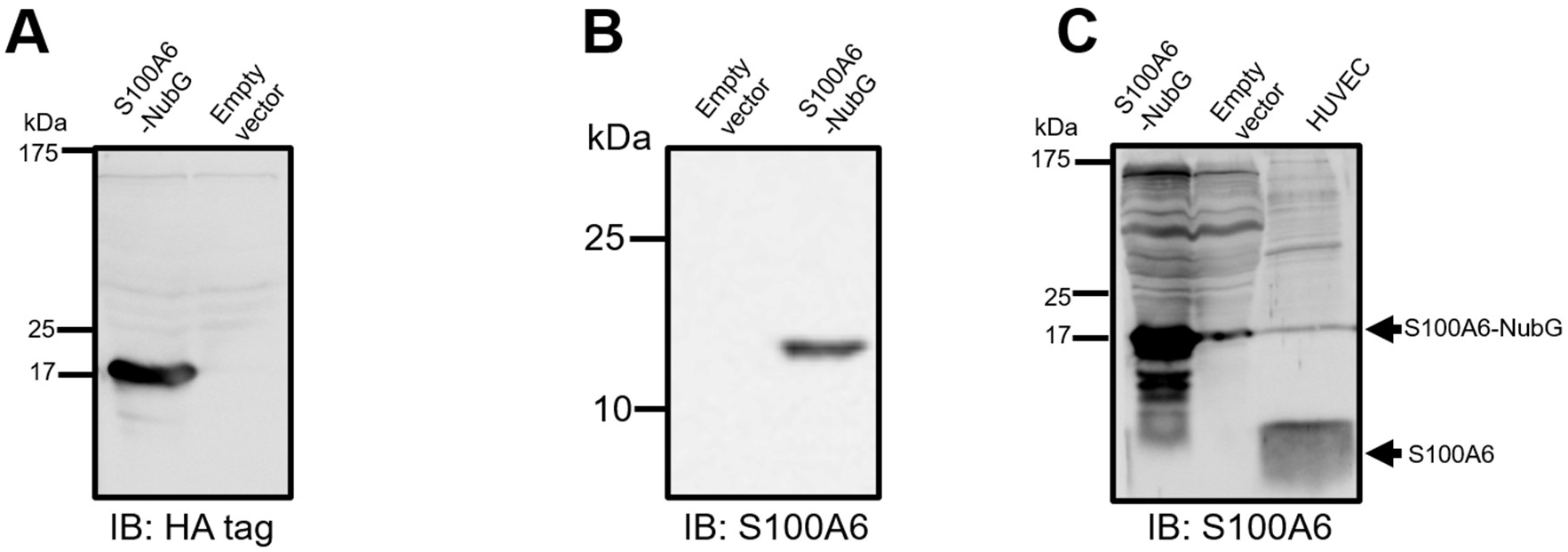
Human S100A6 expression in yeast and primary human endothelial cells. (**A**) Expression and detection of S100A6-NubG in yeast by immunoblotting using anti-HA antibodies. (**D**) Expression and detection of S100A6-NubG in yeast by immunoblotting using anti-S100A6 antibodies. (**E**) Expression and detection of human endothelial S100A6 and S100A6-NubG in yeast by immunoblotting using anti-S100A6 antibodies. Arrowheads denote human endothelial S100A6 and S100A6-NubG fusion proteins respectively.

## References

Apte, R.S., D.S. Chen, and N. Ferrara. 2019. VEGF in Signaling and Disease: Beyond Discovery and Development. Cell. 176:1248–1264.

Autiero, M., J. Waltenberger, D. Communi, A. Kranz, L. Moons, D. Lambrechts, J. Kroll, S. Plaisance, M. De Mol, F. Bono, S. Kliche, G. Fellbrich, K. Ballmer-Hofer, D. Maglione, U. Mayr-Beyrle, M. Dewerchin, S. Dombrowski, D. Stanimirovic, P. Van Hummelen, C. Dehio, D.J. Hicklin, G. Persico, J.M. Herbert, D. Communi, M. Shibuya, D. Collen, E.M. Conway, and P. Carmeliet. 2003. Role of PlGF in the intra- and intermolecular cross talk between the VEGF receptors Flt1 and Flk1. Nat Med. 9:936–943.

Bao, L., A.F. Odell, S.L. Stephen, S.B. Wheatcroft, J.H. Walker, and S. Ponnambalam. 2012. The S100A6 calcium-binding protein regulates endothelial cell-cycle progression and senescence. FEBS J. 279:4576–4588.

Bates, D.O., N. Beazley-Long, A.V. Benest, X. Ye, N. Ved, R.P. Hulse, S. Barratt, M.J. Machado, L.F. Donaldson, S.J. Harper, M. Peiris-Pages, D.J. Tortonese, S. Oltean, and R.R. Foster. 2018. Physiological Role of Vascular Endothelial Growth Factors as Homeostatic Regulators. Compr Physiol. 8:955–979.

Bruns, A.F., S.P. Herbert, A.F. Odell, H.M. Jopling, N.M. Hooper, I.C. Zachary, J.H. Walker, and S. Ponnambalam. 2010. Ligand-stimulated VEGFR2 signaling is regulated by co-ordinated trafficking and proteolysis. Traffic. 11:161–174.

Chehab, T., N.C. Santos, A. Holthenrich, S.N. Koerdt, J. Disse, C. Schuberth, A.R. Nazmi, M. Neeft, H. Koch, K.N.M. Man, S.M. Wojcik, T.F.J. Martin, P. van der Sluijs, N. Brose, and V. Gerke. 2017. A novel Munc13-4/S100A10/annexin A2 complex promotes Weibel-Palade body exocytosis in endothelial cells. Mol Biol Cell. 28:1688–1700.

de Vries, S.J., and A.M. Bonvin. 2011. CPORT: a consensus interface predictor and its performance in prediction-driven docking with HADDOCK. PLoS One. 6:e17695.

Deng, Y., M. Pakdel, B. Blank, E.L. Sundberg, C.G. Burd, and J. von Blume. 2018. Activity of the SPCA1 Calcium Pump Couples Sphingomyelin Synthesis to Sorting of Secretory Proteins in the Trans-Golgi Network. Dev Cell. 47:464–478 e468.

Dominguez, C., R. Boelens, and A.M. Bonvin. 2003. HADDOCK: a protein-protein docking approach based on biochemical or biophysical information. J Am Chem Soc. 125:1731–1737.

Donato, R., G. Sorci, and I. Giambanco. 2017. S100A6 protein: functional roles. Cell Mol Life Sci. 74:2749–2760.

Endres, N.F., T. Barros, A.J. Cantor, and J. Kuriyan. 2014. Emerging concepts in the regulation of the EGF receptor and other receptor tyrosine kinases. Trends Biochem Sci. 39:437–446.

Ewan, L.C., H.M. Jopling, H. Jia, S. Mittar, A. Bagherzadeh, G.J. Howell, J.H. Walker, I.C. Zachary, and S. Ponnambalam. 2006. Intrinsic tyrosine kinase activity is required for vascular endothelial growth factor receptor 2 ubiquitination, sorting and degradation in endothelial cells. Traffic. 7:1270–1282.

Fearnley, G.W., G.A. Smith, I. Abdul-Zani, N. Yuldasheva, N.A. Mughal, S. Homer-Vanniasinkam, M.T. Kearney, I.C. Zachary, D.C. Tomlinson, M.A. Harrison, S.B. Wheatcroft, and S. Ponnambalam. 2016. VEGF-A isoforms program differential VEGFR2 signal transduction, trafficking and proteolysis. Biol Open. 5:571–583.

Fearnley, G.W., G.A. Smith, A.F. Odell, A.M. Latham, S.B. Wheatcroft, M.A. Harrison, D.C. Tomlinson, and S. Ponnambalam. 2014. Vascular endothelial growth factor A-stimulated signaling from endosomes in primary endothelial cells Meth Enzymol. 535:265–292.

Fischer, C., M. Mazzone, B. Jonckx, and P. Carmeliet. 2008. FLT1 and its ligands VEGFB and PlGF: drug targets for anti-angiogenic therapy? Nat Rev Cancer. 8:942–956.

Gampel, A., L. Moss, M.C. Jones, V. Brunton, J.C. Norman, and H. Mellor. 2006. VEGF regulates the mobilization of VEGFR2/KDR from an intracellular endothelial storage compartment. Blood. 108:2624–2631.

Girard, C., N. Tinel, C. Terrenoire, G. Romey, M. Lazdunski, and M. Borsotto. 2002. p11, an annexin II subunit, an auxiliary protein associated with the background K+ channel, TASK-1. EMBO J. 21:4439–4448.

Guo, D., Q. Jia, H.Y. Song, R.S. Warren, and D.B. Donner. 1995. Vascular endothelial cell growth factor promotes tyrosine phosphorylation of mediators of signal transduction that contain SH2 domains. Association with endothelial cell proliferation. J Biol Chem. 270:6729–6733.

Guo, Y., D.W. Sirkis, and R. Schekman. 2014. Protein sorting at the trans-Golgi network. Annu Rev Cell Dev Biol. 30:169–206.

Hubbard, S.J., and J.M. Thornton. 1993. ‘NACCESS’ Computer Program. Department of Biochemistry & Molecular Biology, University College London, London, UK.

Johnsson, N., and A. Varshavsky. 1994. Split ubiquitin as a sensor of protein interactions in vivo. Proc Natl Acad Sci U S A. 91:10340–10344.

Jones, M.C., P.T. Caswell, K. Moran-Jones, M. Roberts, S.T. Barry, A. Gampel, H. Mellor, and J.C. Norman. 2009. VEGFR1 (Flt1) regulates Rab4 recycling to control fibronectin polymerization and endothelial vessel branching. Traffic. 10:754–766.

Jopling, H.M., G.J. Howell, N. Gamper, and S. Ponnambalam. 2011. The VEGFR2 receptor tyrosine kinase undergoes constitutive endosome-to-plasma membrane recycling. Biochem and Biophys Res Comm. 410:170–176.

Koch, S., and L. Claesson-Welsh. 2012. Signal transduction by vascular endothelial growth factor receptors. Cold Spring Harb Perspect Med. 2:a006502.

Kroll, J., and J. Waltenberger. 1997. The vascular endothelial growth factor receptor KDR activates multiple signal transduction pathways in porcine aortic endothelial cells. J Biol Chem. 272:32521–32527.

Lampugnani, M.G., F. Orsenigo, M.C. Gagliani, C. Tacchetti, and E. Dejana. 2006. Vascular endothelial cadherin controls VEGFR-2 internalization and signaling from intracellular compartments. J Cell Biol. 174:593–604.

Lanahan, A., X. Zhang, A. Fantin, Z. Zhuang, F. Rivera-Molina, K. Speichinger, C. Prahst, J. Zhang, Y. Wang, G. Davis, D. Toomre, C. Ruhrberg, and M. Simons. 2013. The neuropilin 1 cytoplasmic domain is required for VEGF-A-dependent arteriogenesis. Dev Cell. 25:156–168.

Larose, L., G. Gish, and T. Pawson. 1995. Construction of an SH2 domain-binding site with mixed specificity. J Biol Chem. 270:3858–3862.

Lee, T.H., S. Seng, M. Sekine, C. Hinton, Y. Fu, H.K. Avraham, and S. Avraham. 2007. Vascular endothelial growth factor mediates intracrine survival in human breast carcinoma cells through internally expressed VEGFR1/FLT1. PLoS Med. 4:e186.

Lemmon, M.A., D.M. Freed, J. Schlessinger, and A. Kiyatkin. 2016. The Dark Side of Cell Signaling: Positive Roles for Negative Regulators. Cell. 164:1172–1184.

Lemmon, M.A., and J. Schlessinger. 2010. Cell signaling by receptor tyrosine kinases. Cell. 141:1117–1134.

Lichtenberger, B.M., P.K. Tan, H. Niederleithner, N. Ferrara, P. Petzelbauer, and M. Sibilia. 2010. Autocrine VEGF signaling synergizes with EGFR in tumor cells to promote epithelial cancer development. Cell. 140:268–279.

Manickam, V., A. Tiwari, J.J. Jung, R. Bhattacharya, A. Goel, D. Mukhopadhyay, and A. Choudhury. 2011. Regulation of vascular endothelial growth factor receptor 2 trafficking and angiogenesis by Golgi localized t-SNARE syntaxin 6. Blood. 117:1425–1435.

Maruyama, I.N. 2015. Activation of transmembrane cell-surface receptors via a common mechanism? The “rotation model”. Bioessays. 37:959–967.

Matsumoto, T., S. Bohman, J. Dixelius, T. Berge, A. Dimberg, P. Magnusson, L. Wang, C. Wikner, J.H. Qi, C. Wernstedt, J. Wu, S. Bruheim, H. Mugishima, D. Mukhopadhyay, A. Spurkland, and L. Claesson-Welsh. 2005. VEGF receptor-2 Y951 signaling and a role for the adapter molecule TSAd in tumor angiogenesis. EMBO J. 24:2342–2353.

McCormack, J.J., M. Lopes da Silva, F. Ferraro, F. Patella, and D.F. Cutler. 2017. Weibel-Palade bodies at a glance. J Cell Sci. 130:3611–3617.

McTigue, M.A., J.A. Wickersham, C. Pinko, R.E. Showalter, C.V. Parast, A. Tempczyk-Russell, M.R. Gehring, B. Mroczkowski, C.C. Kan, J.E. Villafranca, and K. Appelt. 1999. Crystal structure of the kinase domain of human vascular endothelial growth factor receptor 2: a key enzyme in angiogenesis. Structure. 7:319–330.

Mikhaylova, M., P.P. Reddy, and M.R. Kreutz. 2010. Role of neuronal Ca2+-sensor proteins in Golgi-cell-surface membrane traffic. Biochem Soc Trans. 38:177–180.

Mittar, S., C. Ulyatt, G.J. Howell, A.F. Bruns, I. Zachary, J.H. Walker, and S. Ponnambalam. 2009. VEGFR1 receptor tyrosine kinase localization to the Golgi apparatus is calcium-dependent. Exp Cell Res. 315:877–889.

Mundhenk, J., C. Fusi, and M.R. Kreutz. 2019. Caldendrin and Calneurons-EF-Hand CaM-Like Calcium Sensors With Unique Features and Specialized Neuronal Functions. Front Mol Neurosci. 12:16.

Okuse, K., M. Malik-Hall, M.D. Baker, W.Y. Poon, H. Kong, M.V. Chao, and J.N. Wood. 2002. Annexin II light chain regulates sensory neuron-specific sodium channel expression. Nature. 417:653–656.

Otterbein, L.R., J. Kordowska, C. Witte-Hoffmann, C.L. Wang, and R. Dominguez. 2002. Crystal structures of S100A6 in the Ca(2+)-free and Ca(2+)-bound states: the calcium sensor mechanism of S100 proteins revealed at atomic resolution. Structure. 10:557–567.

Pakdel, M., and J. von Blume. 2018. Exploring new routes for secretory protein export from the trans-Golgi network. Mol Biol Cell. 29:235–240.

Rahman, H.N.A., H. Wu, Y. Dong, S. Pasula, A. Wen, Y. Sun, M.L. Brophy, K.L. Tessneer, X. Cai, J. McManus, B. Chang, S. Kwak, N.S. Rahman, W. Xu, C. Fernandes, J.M. McDaniel, L. Xia, L. Smith, R.S. Srinivasan, and H. Chen. 2016. Selective Targeting of a Novel Epsin-VEGFR2 Interaction Promotes VEGF-Mediated Angiogenesis. Circ Res. 118:957–969.

Rezvanpour, A., and G.S. Shaw. 2009. Unique S100 target protein interactions. Gen Physiol Biophys. 28 Spec No Focus:F39–46.

Salikhova, A., L. Wang, A.A. Lanahan, M. Liu, M. Simons, W.P. Leenders, D. Mukhopadhyay, and A. Horowitz. 2008. Vascular endothelial growth factor and semaphorin induce neuropilin-1 endocytosis via separate pathways. Circ Res. 103:e71–79.

Santamaria-Kisiel, L., A.C. Rintala-Dempsey, and G.S. Shaw. 2006. Calcium-dependent and -independent interactions of the S100 protein family. Biochem J. 396:201–214.

Shibuya, M. 2015. VEGF-VEGFR System as a Target for Suppressing Inflammation and other Diseases. Endocr Metab Immune Disord Drug Targets. 15:135–144.

Shibuya, M., and L. Claesson-Welsh. 2006. Signal transduction by VEGF receptors in regulation of angiogenesis and lymphangiogenesis. Exp Cell Res. 312:549–560.

Simons, M., E. Gordon, and L. Claesson-Welsh. 2016. Mechanisms and regulation of endothelial VEGF receptor signalling. Nat Rev Mol Cell Biol. 17:611–625.

Smith, G.A., G.W. Fearnley, I. Abdul-Zani, S.B. Wheatcroft, D.C. Tomlinson, M.A. Harrison, and S. Ponnambalam. 2017. Ubiquitination of basal VEGFR2 regulates signal transduction and endothelial function. Biol Open. 6:1404–1415.

Smith, G.A., G.W. Fearnley, D.C. Tomlinson, M.A. Harrison, and S. Ponnambalam. 2015. The cellular response to vascular endothelial growth factors requires co-ordinated signal transduction, trafficking and proteolysis. Biosci Rep. 35:e00253.

Stagljar, I., C. Korostensky, N. Johnsson, and S. te Heesen. 1998. A genetic system based on split-ubiquitin for the analysis of interactions between membrane proteins in vivo. Proc Natl Acad Sci U S A. 95:5187–5192.

Stoletov, K.V., K.E. Ratcliffe, S.C. Spring, and B.I. Terman. 2001. NCK and PAK participate in the signaling pathway by which vascular endothelial growth factor stimulates the assembly of focal adhesions. J Biol Chem. 276:22748–22755.

Sun, Z., X. Li, S. Massena, S. Kutschera, N. Padhan, L. Gualandi, V. Sundvold-Gjerstad, K. Gustafsson, W.W. Choy, G. Zang, M. Quach, L. Jansson, M. Phillipson, M.R. Abid, A. Spurkland, and L. Claesson-Welsh. 2012. VEGFR2 induces c-Src signaling and vascular permeability in vivo via the adaptor protein TSAd. J Exp Med. 209:1363–1377.

Takahashi, T., S. Yamaguchi, K. Chida, and M. Shibuya. 2001. A single autophosphorylation site on KDR/Flk-1 is essential for VEGF-A-dependent activation of PLC-gamma and DNA synthesis in vascular endothelial cells. EMBO J. 20:2768–2778.

Tatulian, S.A. 2015. Structural Dynamics of Insulin Receptor and Transmembrane Signaling. Biochemistry. 54:5523–5532.

Tiwari, A., J.J. Jung, S.M. Inamdar, D. Nihalani, and A. Choudhury. 2013. The myosin motor Myo1c is required for VEGFR2 delivery to the cell surface and for angiogenic signaling. Am J Physiol Heart Circ Physiol. 304:H687–696.

Vaisman, N., D. Gospodarowicz, and G. Neufeld. 1990. Characterization of the receptors for vascular endothelial growth factor. J Biol Chem. 265:19461–19466.

van Zundert, G.C.P., J. Rodrigues, M. Trellet, C. Schmitz, P.L. Kastritis, E. Karaca, A.S.J. Melquiond, M. van Dijk, S.J. de Vries, and A. Bonvin. 2016. The HADDOCK2.2 Web Server: User-Friendly Integrative Modeling of Biomolecular Complexes. J Mol Biol. 428:720–725.

von Blume, J., A.M. Alleaume, C. Kienzle, A. Carreras-Sureda, M. Valverde, and V. Malhotra. 2012. Cab45 is required for Ca(2+)-dependent secretory cargo sorting at the trans-Golgi network. J Cell Biol. 199:1057–1066.

Yamada, K.H., Y. Nakajima, M. Geyer, K.K. Wary, M. Ushio-Fukai, Y. Komarova, and A.B. Malik. 2014. KIF13B regulates angiogenesis through Golgi to plasma membrane trafficking of VEGFR2. J Cell Sci. 127:4518–4530.

Yang, A.D., E.R. Camp, F. Fan, L. Shen, M.J. Gray, W. Liu, R. Somcio, T.W. Bauer, Y. Wu, D.J. Hicklin, and L.M. Ellis. 2006. Vascular endothelial growth factor receptor-1 activation mediates epithelial to mesenchymal transition in human pancreatic carcinoma cells. Cancer Res. 66:46–51.

Yang, Z., H.M. Kirton, D.A. MacDougall, J.P. Boyle, J. Deuchars, B. Frater, S. Ponnambalam, M.E. Hardy, E. White, S.C. Calaghan, C. Peers, and D.S. Steele. 2015. The Golgi apparatus is a functionally distinct Ca2+ store regulated by the PKA and Epac branches of the beta1-adrenergic signaling pathway. Sci Signal. 8:ra101.

Zhang, Z., K.G. Neiva, M.W. Lingen, L.M. Ellis, and J.E. Nor. 2010. VEGF-dependent tumor angiogenesis requires inverse and reciprocal regulation of VEGFR1 and VEGFR2. Cell Death Differ. 17:499–512.

Zhou, H.J., Z. Xu, Z. Wang, H. Zhang, M. Simons, and W. Min. 2018. SUMOylation of VEGFR2 regulates its intracellular trafficking and pathological angiogenesis. Nat Commun. 9:3303.

